# A Plant Protein NbP3IP Induces Autophagy and Mediates the Autophagic Degradation of RSV p3 to Inhibit Viral Infection

**DOI:** 10.1101/532275

**Authors:** Liangliang Jiang, Yuwen Lu, Xiyin Zheng, Xue Yang, Ying Chen, Tianhao Zhang, Xing Zhao, Shu Wang, Xia Zhao, Xijiao Song, Xiangxiang Zhang, Jiejun Peng, Hongying Zheng, Lin Lin, Stuart MacFarlane, Jianping Chen, Yule Liu, Fei Yan

## Abstract

In plants, autophagy is involved in responses to viral infection. However, understanding of new host factors mediating autophagic clearance of plant viruses is very limited. We here identified a new host factor NbP3IP participating in autophagy-mediated plant defense against viral infection. NbP3IP interacted with p3, a RNA silencing suppressor encoded by Rice stripe virus (RSV), a negative-strand RNA virus, and mediated its autophagic degradation. NbP3IP could also interact with NbATG8f, which was required for NbP3IP-miediated p3 degradation. Overexpression of NbP3IP induced autophagy and down-regulation of NbP3IP reduced autophagy. Both overexpression of NbP3IP and silencing of GAPC, which also induces autophagy, inhibited RSV infection. In contrast, silencing of ATG7 promoted RSV infection. Thus, through identification of a new potential selective autophagy receptor P3IP, we revealed a new mechanism of autophagy-mediated plant defense against plant viruses and provided the first evidence that plant autophagy can also play an antiviral role against negative-strand RNA viruses.

## INTRODUCTION

Rice stripe virus (RSV), transmitted by the small brown planthopper (SBPH; *Laodelphax striatellus* Fallén), causes serious epidemics in East Asia, including China, Japan and Korea (Cheng et al., 2008). RSV belongs to the genus *Tenuivirus* and only infects plants of the family *Poaceae* by natural SBPH transmission, which is a barrier to studies exploring the pathogenesis of RSV and its interaction with plants in the field or with its insect vector. In the laboratory, however, RSV can infect the experimental plant *Nicotiana benthamiana* by mechanical inoculation, which has been adopted as a very useful model system for studying RSV-plant interactions (Xiong et al., 2008; Yuan et al., 2011; Zhang et al., 2012; Kong et al., 2013; Fu et al., 2018). RSV has four single-stranded RNA genome segments. RNA1 (∼9 kb) is negative-sense and has a single open reading frame (ORF), encoding the RNA-dependent RNA polymerase (RdRP), in its complementary strand. Each of the other three segments (RNA2, 3.5 kb; RNA3, 2.5 kb; RNA4, 2.2 kb) are ambisense and contain two non-overlapping ORFs on opposite strands, separated by a non-coding intergenic region (IR) that functions in termination of transcription (Zhu et al., 1991; Zhu et al., 1992; Hamamatsu et al., 1993; Takahashi et al., 1993; Qu et al., 1997; Wu et al., 2013). RNA2 encodes two proteins, p2 and pc2. p2 has weak RNA silencing suppressor activity, and pc2 is a nonstructural protein with unknown function, but which is processed in insect cells into pc2-N and pc2-C (Du et al., 2011; Zhao et al., 2012). On RNA3, p3 and pc3 are, respectively, the primary viral suppressor of RNA silencing (VSR) and the nucleocapsid protein (Xiong et al., 2009). RNA4 encodes a disease-specific protein p4 that can interact with the plant PsbP protein causing viral symptoms, and also with the viral movement protein pc4 (Xiong et al., 2008; Yuan et al., 2011; Zhang et al., 2012; Kong et al., 2013). RSV interferes with s-acylation of remorin and induces its autophagic degradation to facilitate RSV infection (Fu et al., 2018).

Autophagy is a conserved eukaryotic mechanism that mediates the degradation of cytoplasmic components and damaged organelles through lysosomal pathways. This process involves multiple autophagy-related (ATG) proteins. Autophagy can be induced by starvation, oxidative stress, drought, salt and pathogen invasion in plants (Bassham, 2007; Liu et al., 2009; Han et al., 2011; Hayward and Dinesh-Kumar, 2011; Bozhkov, 2018). In plants, increasing evidence shows that autophagy plays an important role in defense against viral infection (Hafren et al., 2017; Haxim et al., 2017; Li et al., 2017; Hafren et al., 2018; Li et al., 2018; Yang et al., 2018). For example, it has been found that Cotton leaf curl Multan virus (CLCuMuV) infection can induce autophagy in *N. benthamiana* (Haxim et al., 2017). NbATG8f, a key factor in autophagy, could interact with βC1, the VSR encoded by the satellite DNA of CLCuMuV, which directs βC1 to autophagosomes for degradation and hence inhibits the infection of CLCuMuV. Beclin1, one of the central ATGs that is upregulated in Turnip mosaic virus-infected plants, was found to interact with the viral RdRP, NIb, and mediated its degradation (Li et al., 2018). Beclin1-mediated NIb degradation was inhibited by autophagy inhibitors and, moreover, deficiency of Beclin1 or ATG8a enhanced NIb accumulation and promoted viral infection, indicating that Beclin1-mediated autophagic degradation of NIb plays a defensive role against TuMV infection. Autophagy was also shown to inhibit the infection of another positive-sense RNA virus, Barley stripe mosaic virus (BSMV), in *N. benthamiana* (Yang et al., 2018). Moreover, the BSMV γb protein was found to suppress the autophagy process by disrupting the interaction between ATG7 and ATG8 (Yang et al., 2018).

Here, we describe a new mechanism of autophagy-mediated plant defense against plant viruses through a new selective autophagy receptor P3IP. Additionally, our work is the first to demonstrate the role of autophagy in plant negative-strand virus infection. We show that the VSR of RSV, p3, interacts with a previously unknown protein of *N. benthamiana* (NbP3IP). We show that NbP3IP also interacts with ATG8f, and demonstrate that this interaction is responsible for the autophagic degradation of p3. We further demonstrate that the overexpression of NbP3IP induces autophagy, while silencing of the *NbP3IP* gene inhibits autophagy, confirming its role in the regulation of autophagy in plants. Our discovery of the role of NbP3IP in autophagy offers new opportunities to understand the mechanisms that underlie this essential cellular process.

## RESULTS

### The plant protein NbP3IP interacts with RSV p3

To identify possible host proteins that interact with RSV p3, a cDNA library of *N. benthamiana*, the experimental host of RSV, was used for a yeast two-hybrid protein screening with p3 as the bait. One of the candidate proteins designated as <u>p3</u> interacting <u>p</u>rotein in *<u>N</u>. <u>b</u>enthamiana* (NbP3IP) was selected for further analysis. Sequencing analysis showed that this cDNA clone contained an ORF of 480 nucleotides that was predicted to encode an 18-kDa protein. In plants, the expression level of the *NbP3IP* gene was not altered significantly at 9 days post infection (dpi) with RSV but was increased at 20 dpi (Supplementary Fig. S1A). The subcellular localization of NbP3IP was examined by transiently expressing the protein with GFP fused to its C-terminus in *N. benthamiana* epidermal cells, which showed that NbP3IP was localized to cytoplasm and nucleus (Supplementary Fig. S1B).

Further Y2H tests confirmed the strong and specific interaction of NbP3IP with RSV p3 (Fig. 1A). To corroborate that this interaction also occurred in the plant, bimolecular fluorescence complementation (BiFC) and co-immunoprecipitation (Co-IP) experiments were performed. We cloned the *NbP3IP* and RSV *p3* genes into the split-YFP vectors pCV-YFPn-C and pCV-YFPc-C (Lu et al., 2011; Jiang et al., 2014) to express NbP3IP and p3 with an N-terminal split-YFP tag in *N. benthamiana* plants. Co-expression of NbP3IP and p3 produced YFP fluorescence at 3 dpi confirming an interaction between these two proteins, whereas, co-expression of NbP3IP with a control protein (GUS) did not (Fig. 1B). Similar BiFC experiments were done to co-express NbP3IP with other viral silencing suppressor proteins (potato virus X (PVX) p25; tomato bushy stunt virus (TBSV) p19; turnip mosaic virus (TuMV) HC-Pro). None of these proteins was found to interact with NbP3IP (Supplementary Fig. S2). In a Co-IP assay, plasmids encoding NbP3IP-eGFP (C-terminal GFP tag) and p3-Myc (C-terminal Myc tag) were transiently expressed in *N. benthamiana*, with pc3-Myc (RSV complementary strand ORF) used as a non-interacting (negative) control. Leaf lysates were immunoprecipitated with anti-GFP beads and any co-precipitated protein detected using an anti-Myc antibody. These experiments demonstrated that NbP3IP interacted with p3 *in vivo* (Fig. 1C). A series of deletion mutants of p3 were created and tested for interaction with NbP3IP in a BiFC assay. The various deletions encompassed four helical regions (H1 to H4) that were previously shown to be important for p3 self-interaction (Kim et al., 2017) and a positively-charged region (NLS) that was shown to be involved in nuclear localization and silencing suppression activity (Xiong et al., 2009) (Fig. 1D). Using this approach, the p3 truncated fragments F6, F7 and F8 were unable to interact with NbP3IP (Fig. 1E). Thus, the NbP3IP-interacting regions of RSV p3 were mapped to the N-terminal region (F4) and C-terminal region (F5) and excluded the H1-H4 and NLS domains.

**Fig.1.**
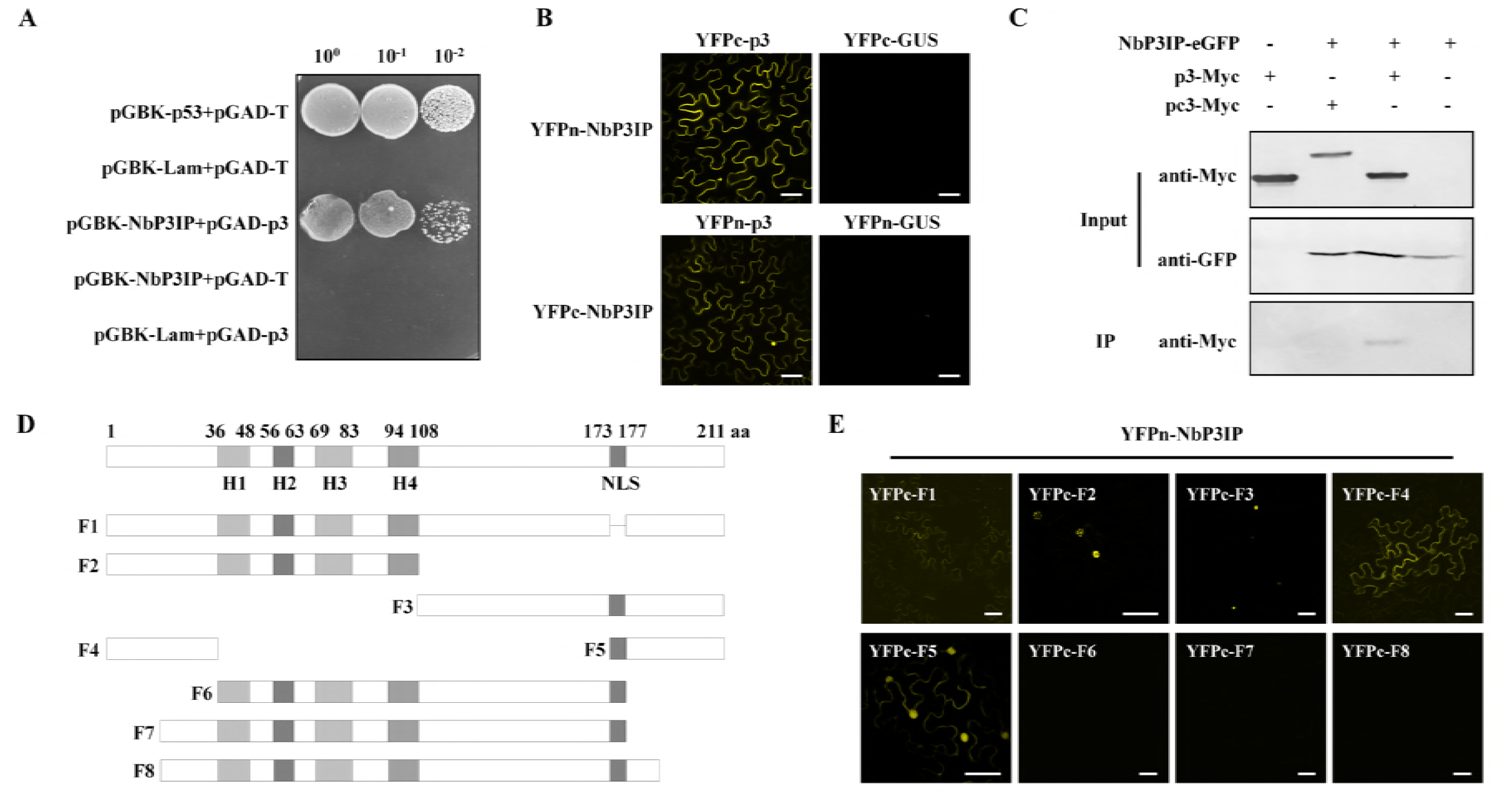
NbP3IP interacts with RSV p3. (A) Yeast two-hybrid assay demonstrates interaction between NbP3IP and p3. NbP3IP was fused with the GAL4 binding domain (pGBK-NbP3IP) and p3 was fused with the GAL4 activation domain (pGAD-p3). Yeast co-transformed with pGBK-53 + pGAD-T served as a positive control, and yeast co-transformed with vectors pGBK-Lam + pGAD-T, pGBK-NbP3IP + pGAD-T or pGBK-Lam + pGAD-p3 served as negative controls. Serial 10-fold dilutions of co-transformed yeast cells were plated on synthetic defined (SD) medium lacking tryptophan, leucine, histidine and adenine and colony growth indicated interaction between the paired proteins. (B) BiFC assay demonstrating interaction between NbP3IP and p3 in the leaves of *N. benthamiana* at 60 hours post infiltration (hpi). The N- or C-terminal fragments of YFP were fused to the N-terminus of NbP3IP and p3. The GUS protein was used as a non-interacting, negative control. Bars, 50 μm. (C) Co-immunoprecipitation (Co-IP) analysis of NbP3IP-eGFP and p3-Myc expressed in *N. benthamiana* leaves by agroinfiltration. Separate expression of NbP3IP-eGFP and p3-Myc, and co-expression of NbP3IP-eGFP and pc3-Myc were used as negative controls. (D) A schematic diagram of the wild-type p3 protein and 8 truncated p3 constructs (F1 to F8). The mutations include a NLS deletion mutant (F1), a C-terminal deletion mutant (F2), an N-terminal deletion mutant (F3), 1-36 aa of p3 (F4), 173-211 aa of p3 (F5), 37-177 aa of p3 (F6), 21-177 aa of p3 (F7), 21-190 aa of p3 (F8). H1 to H4 represent four helical regions. (E) BiFC assays between NbP3IP and each of the 8 mutants of p3 in leaves of *N. benthamiana* at 60 hpi. Bars, 50 μm.

### NbP3IP inhibits the local VSR ability of RSV p3

Initially, the VSR ability of p3 was confirmed using a co-infiltration assay in GFP-expressing (16c) transgenic plants. These plants were infiltrated with an agrobacterium culture expressing GFP (pBIN-GFP) that initially increases GFP levels but then initiates post-transcriptional gene silencing to degrade GFP mRNAs and abolish GFP fluorescence. Simultaneously the plants were also infiltrated with agrobacterium expressing either RSV p3 with a Myc tag (p3-Myc), GUSp-Myc (5’ terminal 500bp of the β-glucuronidase gene) or NbP3IP-Myc. At 5 days post infiltration (dpi), green fluorescence was almost absent in the patch containing pBIN-GFP only, showing that silencing of the GFP gene had occurred (Fig 2A). In addition, there was little or no GFP in the patches containing both pBIN-GFP and GUSp-Myc or NbP3IP-Myc, showing that these proteins could not suppress silencing of the GFP gene. However, strong GFP fluorescence was apparent in the patch containing pBIN-GFP and p3-Myc, confirming that the RSV p3 protein does suppress RNA silencing in this assay system. The increased level of GFP in the p3-treated patch was further confirmed by western blotting, as was the expression of GUSp-Myc, NbP3IP-Myc and p3-Myc in these plants (Fig. 2B).

**Fig.2.**
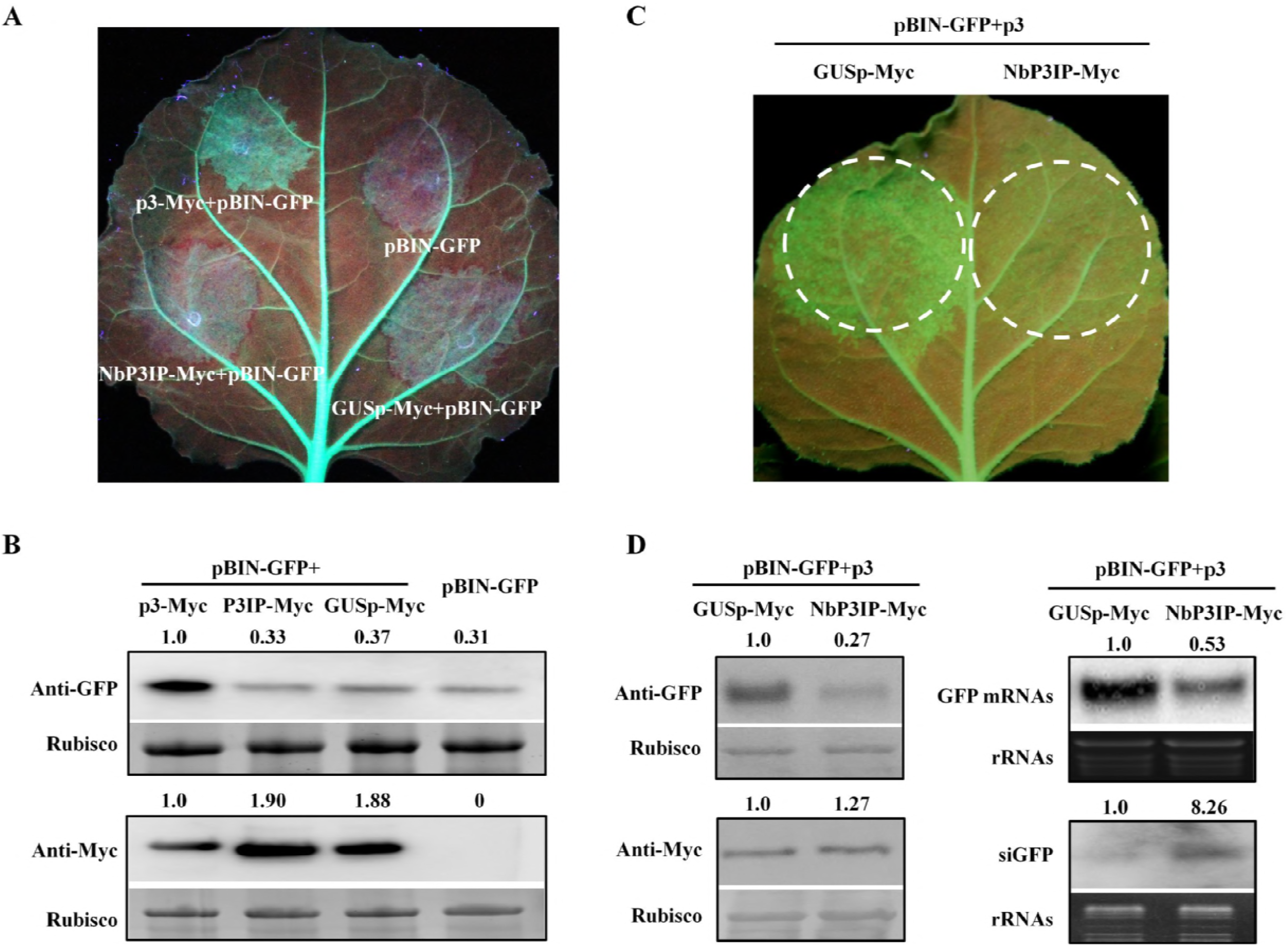
The expression of NbP3IP inhibits the local silencing suppression activity of RSV p3. (A) Expression of pBIN-GFP promotes RNA silencing of the GFP gene in line 16c plants (top, right patch) but co-expression with p3-Myc suppresses RNA silencing and promotes GFP expression (top, left patch). The expression of NbP3IP-Myc or GUSp-Myc (1-166 aa of GUS) did not suppress RNA silencing initiated by pBIN-GFP (bottom patches, left and right). The plants were photographed under UV light at 5 dpi. (B) The accumulation of GFP in the infiltrated patches was detected by western blotting. The expression of p3-Myc, NbP3IP-Myc and GUSp-Myc protein were detected by western blotting with an anti-Myc antibody. The GFP protein levels were normalized to rubisco, and the level in each patch calculated in relation to the p3-Myc sample. (C) The silencing suppression activity of p3 was inhibited by the co-expression of NbP3IP in pBIN-GFP-infiltrated 16c transgenic plants (right side), whereas, co-expression of GUSp-Myc with pBIN-GFP did not affect p3-mediated silencing suppression (left side). The plants were photographed under UV light at 5 dpi. (D) The accumulations of GFP protein, GFP mRNA and GFP siRNAs were detected by western blot and northern blot. The expression of GUSp-Myc and NbP3IP-Myc protein were detected by western blot with Myc antibody. The GFP protein levels were normalized to rubisco, and the relative levels calculated in relation to the GUSp-Myc sample. The GFP mRNA and siRNA levels were normalized to rRNA, and the relative levels calculated in relation to the GUSp-Myc sample.

In a different experiment 16c plants were co-infiltrated with a mixture of pBIN-GFP, p3 and either GUSp-Myc or NbP3IP-Myc. Suppression of GFP silencing by p3 continued in the presence of GUSp-Myc (Fig. 2C, left) but was prevented in the presence of NbP3IP-Myc (Fig, 2C, right). Western blotting of GFP proteins extracted from the infiltrated patches confirmed that the accumulation of GFP in the combination of pBIN-GFP and p3 plus GUSp was higher than in that of pBIN-GFP and p3 plus NbP3IP (Fig. 2D). Northern blotting of GFP mRNA from the patches revealed that the expression level of GFP mRNA in the GUSp-infiltrated patch was higher than the NbP3IP-infiltrated patch and, correspondingly, the level of GFP-derived siRNA was higher in the NbP3IP-containing patch compared with the GUSp control patch (Fig. 2D). These results indicated that the co-expression of NbP3IP inhibited the RNA silencing suppressor function of p3.

To examine whether NbP3IP interferes with silencing suppression by a different VSR, the GFP patch assay was repeated in the presence of the PVX p25 protein, the TBSV p19 protein or the TuMV HC-Pro. Here, at 5 dpi, all three VSRs suppressed silencing of GFP regardless of whether they were co-expressed with GUSp or with NbP3IP (Supplementary Fig. S3). Thus, NbP3IP was shown to specifically interfere with silencing suppression activity of RSV p3 only.

### NbP3IP mediates the degradation of p3 via the autophagy pathway

It has been reported that deletion of the p3 nuclear localization signal significantly reduces its silencing suppression activity (Xiong et al., 2009). To investigate whether NbP3IP influences the function of the p3 silencing suppressor by altering its subcellular localization, p3-GFP (C-terminal GFP tag) and NbP3IP were co-expressed in *N.benthamiana*, with unfused RFP (transiently expressed from co-infiltrated pCV-RFP) being used as a reference protein to allow comparisons of expression levels between treatments. As a further control, p3-GFP was co-expressed with GUSp and unfused RFP. Confocal microscopy showed that p3 was localized in both the nucleus and cytoplasm as previously reported (Xiong et al., 2009). However, the intensity of p3-GFP fluorescence was much lower when co-expressed with NbP3IP than when co-expressed with GUSp, but no intensity change was seen for the internal control RFP protein (Fig. 3A). Western blotting using an anti-GFP antibody confirmed that p3 accumulated to a much lower level when co-expressed with NbP3IP (Fig. 3A). Semi-quantitative RT-PCR testing showed that the levels of p3-GFP mRNAs were similar in both the NbP3IP- and GUSp-treated plants, suggesting that the reduced p3 accumulation could be related to a lower stability or higher turnover rate of the protein (Fig. 3A). In a control experiment, co-expression with NbP3IP did not change the subcellular location or the accumulation level of unfused GFP protein (Supplementary Fig. S4A, S4B), indicating that NbP3IP specifically affected the accumulation of RSV p3. In contrast, the subcellular localization of NbP3IP-GFP was not affected by co-expression with RSV p3, or any of the other non-interacting silencing suppressor proteins (p25, p19 or HC-Pro) (Supplementary Fig. S4C).

**Fig.3.**
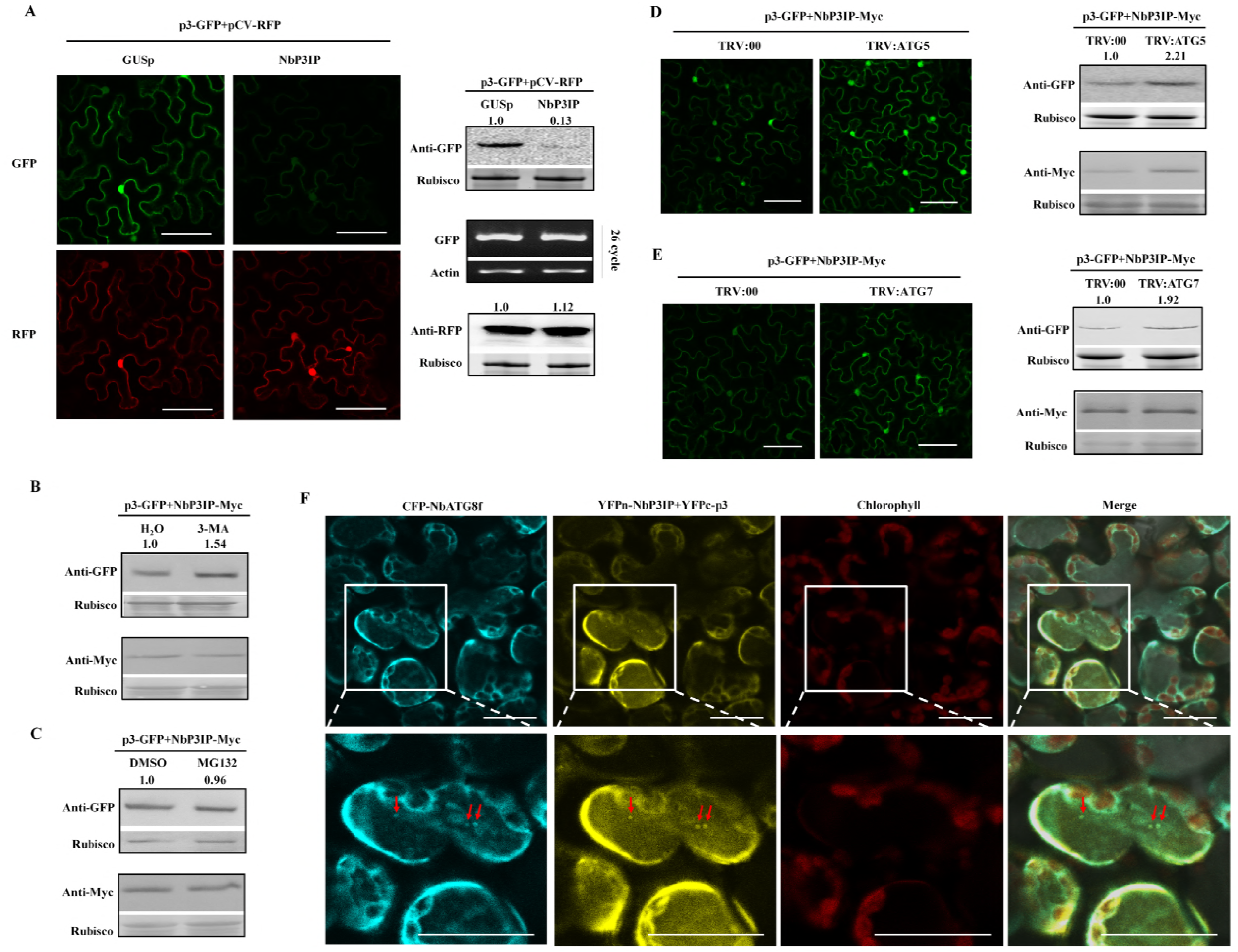
NbP3IP mediates the degradation of p3 via the autophagy pathway. (A) Protein accumulation analysis of p3-GFP (empty RFP as an expressing reference) co-expressed with NbP3IP or GUSp. p3-GFP fluorescence was detected by confocal microscopy at 60 hpi. Bars, 20 μm. The accumulation of the p3 protein was analyzed by western blotting with an anti-GFP antibody. The level of p3-GFP co-expressed with GUSp was normalized to rubisco, and used as the reference for relative expression calculations. Semi-quantitative RT-PCR was used for analysis of p3-GFP transcripts. Actin served as an internal amplification standard. The accumulation of RFP was detected by western blotting with anti-RFP antibody. The level of RFP was normalized to rubisco. (B) The autophagy inhibitor 3-MA reduces NbP3IP-mediated degradation of p3-GFP. p3-GFP and NbP3IP-Myc were co-expressed in *N. benthamiana* leaves for 48 h, followed by a H_2_O or 10 mM 3-MA treatment for 16 h. The value represents protein accumulation relative to rubisco, and the relative levels were calculated in relation to the H_2_O treatment. (C) The 26S Proteasome inhibitor MG132 has no effect on the NbP3IP-mediated degradation of p3-GFP. Co-expressed p3-GFP and NbP3IP-Myc in *N. benthamiana* leaves for 48 h, followed by DMSO or MG132 treatment for 16 h. The value represents protein accumulation relative to rubisco, and the relative levels were calculated in relation to the DMSO treatment. (D) Silencing of *NbATG5* inhibited NbP3IP-mediated degradation of p3. Confocal micrographs show cells co-expressing p3-GFP and NbP3IP-Myc in plants with *NbATG5* silencing (TRV:ATG5) or with a non-silenced control (TRV:00) at 60 hpi. Bars, 25 μm. The accumulation of NbP3IP-Myc and p3-GFP were analyzed by western blotting. The value represents protein accumulation relative to rubisco, and the relative levels were calculated in relation to the TRV:00 treatment. (E) Silencing of *NbATG7* inhibited NbP3IP-mediated degradation of p3. Confocal micrographs shows cells co-expressing p3-GFP and NbP3IP-Myc in plants with *NbATG7* silencing (TRV:ATG7) or with a non-silenced control (TRV:00) at 60 hpi. Bars, 25 μm. Protein analysis was done as for panel D. (F) The NbP3IP and p3 interaction complex co-localized with CFP-NbATG8f in *N. benthamiana* leaf epidermal cells. Confocal imaging was done at 60 hpi. Yellow fluorescence indicates co-localized interaction of NbP3IP and p3. Cyan fluorescence indicates the location of NbATG8f. Typical autophagic structures (cyan; red arrows) and the NbP3IP-p3 complex (yellow; red arrows) were observed to co-localize in the cytoplasm. Bars, 25 μm.

To elucidate which pathway was responsible for the degradation of the p3 protein when co-expressed with NbP3IP, further infiltrations were done in the presence of either MG132 (an inhibitor of the 26S proteasome pathway) or 3-methyladenine (3-MA) (an inhibitor of autophagy) (Seglen and Gordon, 1982; Tanida et al., 2005). Western blotting showed that accumulation of p3-GFP when co-expressed with NbP3IP was increased by 3-MA treatment (Fig. 3B). but not by MG132 treatment (Fig. 3C). Also, the autophagy inhibitor 3-MA had no effect on accumulation of p3-GFP in the absence of NbP3IP (Supplementary Fig. S4D). To further show that NbP3IP-mediated degradation of p3 operates through the autophagy pathway, a TRV-based virus induced gene silencing (VIGS) experiment was done to silence two autophagy related genes *NbATG5* and *NbATG7* (Supplementary Fig. S5A). Silencing either *NbATG5* or *NbATG7* inhibited NbP3IP-mediated degradation of p3 (Fig. 3D and 3E).

It is known that ATG8 family proteins execute important functions during autophagy in various species (Kabeya et al., 2000; Yoshimoto et al., 2004; Nakatogawa et al., 2007; Xie et al., 2008). Using a previously validated approach (Haxim et al., 2017), we expressed Cyan Fluorescent Protein (CFP)-tagged *N. benthamiana* ATG8f (CFP-NbATG8f) as an autophagosome marker and examined whether it co-localized in the cell with the interacting complex of p3 and NbP3IP. As shown in Figure 3F, the yellow fluorescence of the YFPn-NbP3IP/YFPc-p3 BiFC pair was spatially associated with the Cyan fluorescence of CFP-NbATG8f, including the overlapping punctate structures in the cytoplasm and marked by red arrows (Fig. 3F). Combining all of the above results, we suggest that the degradation of the p3 protein induced by NbP3IP depends on the autophagy pathway, rather than the proteasome pathway.

### Degradation of p3 via autophagy depends on its interaction with NbP3IP

Down-regulation of cytosolic glyceraldehyde-3-phosphate dehydrogenase (GAPCs) gene expression by VIGS significantly activates autophagy in *N.benthamiana*, which can be visualized by the increase in appearance in the cytoplasm of autophagic bodies containing the NbATG8f protein (Han et al., 2015). Hence, we employed this strategy to induce autophagy and tested whether the degradation of p3 via the autophagy pathway occurred in the absence of NbP3IP. Knockdown of GAPC1, GAPC2 and GAPC3, inducing autophagy, was confirmed by qRT-PCR and confocal microscopy (Supplementary Fig. S5B, S5C, S5D). As shown in Figure 4A, the accumulation of expressed p3-GFP in GAPCs-silenced plants (TRV:GAPCs) showed no obvious differences compared with control (TRV:00) plants. However, the accumulation of p3-GFP co-expressed with NbP3IP in TRV:GAPCs plants was evidently lower than when it was co-expressed with NbP3IP in TRV:00 plants (Fig. 4B). Thus, even when autophagy is stimulated, RSV p3 is not degraded unless it is co-expressed with NbP3IP. The previous experiments showed that mutant F8 of p3 did not interact with NbP3IP. As shown in Figure 4C, in TRV:GAPCs plants, F8-GFP was not degraded via autophagy even when co-expressed with NbP3IP, suggesting that the autophagic degradation of p3 required its physical interaction with NbP3IP.

**Fig.4.**
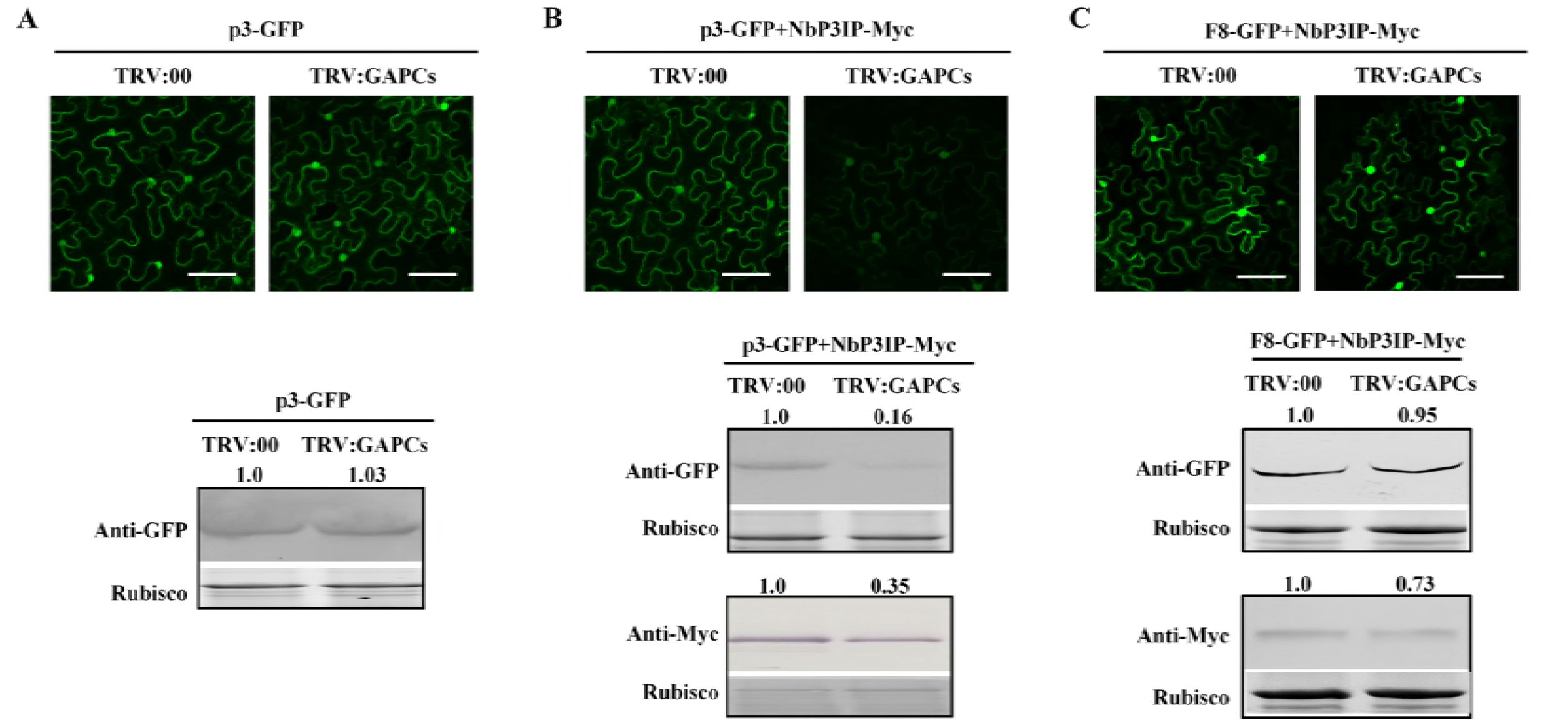
Degradation of p3 via autophagy depends on its interaction with NbP3IP. (A) Activating autophagy by GAPCs silencing has no effect on the accumulation of p3 when expressed alone. Confocal micrographs show cells expressing p3-GFP in plants with *GAPCs* silencing (TRV:GAPCs) or a non-silenced control (TRV:00) at 60 hpi. Bars, 25 μm. Western blot shows the accumulation of p3-GFP in these plants. (B) Analysis of p3 accumulation in *GAPCs* silenced plants or non-silenced plants when co-expressed with NbP3IP. Confocal micrographs shows cells co-expressing p3-GFP and NbP3IP-Myc in *GAPCs* silenced (TRV:GAPCs) or non-silenced control (TRV:00) plants at 60 hpi. Bars, 25 μm. Western blot shows the accumulation of p3-GFP and NbP3IP-Myc in these plants. (C) The p3 deletion mutant F8 that does not interact with NbP3IP is not degraded by the autophagy pathway. Confocal micrographs shows cells co-expressing F8-GFP and NbP3IP-Myc in plants with *GAPCs* silencing (TRV:GAPCs) or a non-silenced control (TRV:00) at 60 hpi. Bars, 25 μm. Western blot shows the accumulation of p3-GFP and NbP3IP-Myc in these plants.

### NbP3IP mediates p3 degradation by interacting with NbATG8f

It has been reported that ATG8 family proteins act as the cargo acceptor participating in the degradation of several plant virus proteins (Haxim et al., 2017; Li et al., 2018). To determine whether p3 or NbP3IP interacted with NbATG8, BiFC was employed in a paired interaction assay. As shown in Figure 5A (upper panels), co-expression of YFPn-NbP3IP and YFPc-NbATG8f produced YFP fluorescence in the cytoplasm and nucleus indicating an interaction, whereas, YFPc-NbATG8f and YFPn-p3 did not interact. In a Co-IP assay using GFP-TRAP beads, NbP3IP-Myc was specifically co-precipitated by GFP-NbATG8f (Fig. 5B), confirming the interaction between NbP3IP and NbATG8f.

**Fig. 5.**
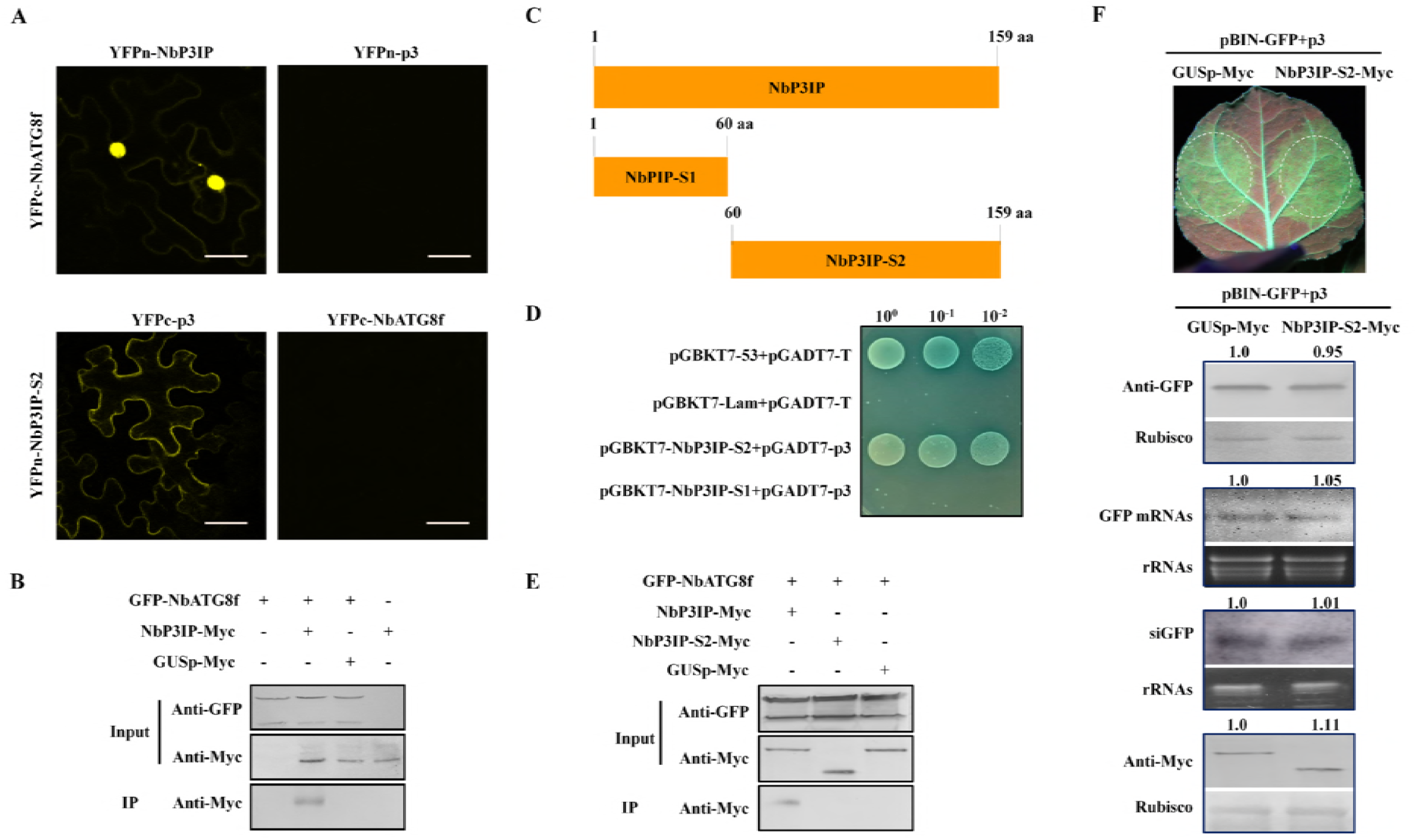
NbP3IP mediates p3 degradation by interacting with NbATG8f. (A) BiFC assay demonstrates interaction between NbP3IP and NbATG8f (upper left panel) and between p3 and C-terminal S2 fragment of NbP3IP (lower left panel). No interaction detected between RSV p3 and NbATG8f (upper right panel) or between NbP3IP S2 fragment and NbATG8f (lower right panel). Bars, 25μm. (B) Co-IP analysis of GFP-NbATG8f and NbP3IP-Myc expressed in *N. benthamiana* leaves by agroinfiltration. NbP3IP-Myc is precipitated only when in the presence of NbATG8f. (C) The diagram shows the two truncated mutants of NbP3IP used in this work. (D) Yeast two-hybrid assays to show interaction between NbP3IP-S2 and RSV p3 but not between NbP3IP-S1 and RSV p3. BK-53 and AD-T are known to interact, whereas BK-Lam and AD-T do not interact. Serial 10-fold dilutions of yeast cells were plated on synthetic defined (SD)/-Trp, -Leu, -His, -Ade medium. (E) Co-IP analysis shows that GFP-NbATG8f interacts with NbP3IP-Myc but not NbPIP-S2-Myc. GUSp-Myc is used as a non-interacting control protein. (F) Suppression of pBIN-GFP-induced silencing of GFP by co-expression of p3 is not inhibited by NbPIP-S2-Myc. GUSp-Myc was used as a non-inhibiting control. The plants were photographed under UV light at 5 dpi. The accumulations of GFP protein and mRNA were detected by western blot and northern blot. The accumulation of GFP-derived siRNA was analyzed in small RNA northern blot assay. The expression of GUSp-Myc and NbP3IP-Myc protein were detected by western blot with Myc antibody. The accumulations of GFP protein, GFP mRNA and GFP siRNAs were detected by western blot and northern blot. The expression of GUSp-Myc and NbP3IP-S2-Myc protein were detected by western blot with Myc antibody. The GFP and Myc-tagged protein levels were normalized to rubisco, and the relative levels calculated in relation to the GUSp-Myc sample. The GFP mRNA and siRNA levels were normalized to rRNA, and the relative levels calculated in relation to the GUSp-Myc sample

To further investigate NbP3IP, two mutant constructs of NbP3IP were made containing either only the N-terminal amino acids 1-60 (mutant NbP3IP-S1) or the C-terminal amino acids 60-159 (mutant NbP3IP-S2) (Fig. 5C). In contrast to wild-type NbP3IP (Fig. 5B) the NbP3IP-S2 fragment did not co-precipitate with NbATG8f (Fig. 5E), although, it did interact with RSV p3 in Y2H (Fig. 5D) and BiFC assays (Fig. 5A, lower left panels). Nevertheless, even though mutant NbP3IP-S2 could interact with p3, this interaction did not inhibit the VSR ability of p3 (Fig. 5F), suggesting that NbP3IP is required to interact with NbATG8f for the degradation of RSV p3 to occur.

### Autophagy is induced during RSV infection and negatively regulates its infection

Due to the involvement of NbATG8f in the degradation of p3, we investigated whether RSV infection had any effect on autophagy. We applied quantitative real-time reverse-transcription PCR (qRT-PCR) to examine the transcription levels of various autophagy-related genes during RSV infection. As shown in figure 6A, mRNA levels of *NbATG2*, *NbATG4*, *NbATG6*, *NbATG7*, *NbATG9*, *NbPI3K* and *NbVps15* were all upregulated in RSV-infected upper leaves at 20 dpi. We also used CFP-NbATG8f as an autophagosome marker to observe the increased number of punctate autophagic bodies in RSV infected plants compared to those in uninfected plants (Fig. 6B). These results indicated that RSV infection activated autophagy.

Since RSV p3 was degraded via the autophagy pathway, we hypothesized that autophagy negatively regulates RSV infection. To validate this hypothesis, TRV:GAPCs plants, in which autophagy was activated (described above; Supplementary Fig. S5C, S5D), were infected with RSV. Compared to control plants (TRV:00), the upper leaves of TRV:GAPCs plants showed less leaf curling, a known symptom of RSV infection (Fig. 6C) and contained lower levels of virus as revealed by analysis of accumulation of RSV capsid protein mRNA and protein (Fig. 6D). Meanwhile, in ATG7-silenced and ATG5-silenced plants (TRV:ATG7, TRV:ATG5), in which the autophagy activity was inhibited, RSV infection symptoms were more severe (increased mosaic and dwarfing) and were accompanied by higher accumulation levels of RSV capsid protein mRNA and protein (Fig. 6E, 6F and Supplementary Fig. S5E). These results indicated that autophagy operates to reduce RSV infection in plants.

**Fig.6.**
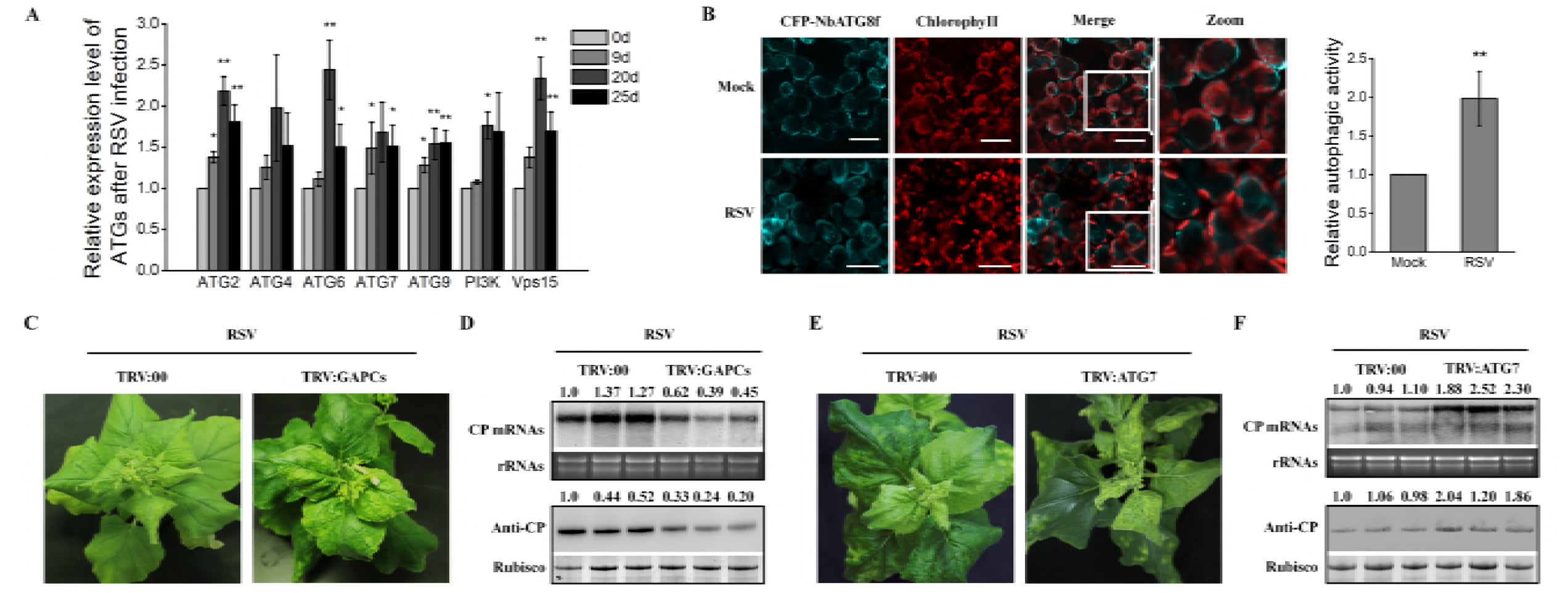
RSV infection induces autophagy and is negatively affected by autophagy. (A) Expression of *ATGs* is up-regulated after RSV infection. Relative quantification of mRNA levels of autophagy-related genes (ATGs) in RSV infected plants at 0, 9, 20, 25 dpi. Values represent means ± SD from three independent experiments. Single asterisk indicates *P*<0.05, double asterisks indicate *P*<0.01 of significant difference between RSV infected 0 day plants and other plants (Student’s *t*-test, two-sided). (B) Confocal micrographs showing *N. benthamiana* leaf cells, infected with RSV at 20 dpi, expressing CFP-NbATG8f at 60 hpi. Autophagosomes and autophagic bodies are revealed as CFP positive puncta in mesophyll cells. Chloroplast autofluorescence is in red. Bars, 10 μm. Relative autophagic activity in RSV infected plants compared to mock inoculated plants. Autophagic bodies were counted from approximately 150 cells for each treatment in three independent experiments. Values represent the means ± SD. Double asterisks indicate P<0.01 of significant difference between control plants and RSV infected plants (Student’s t-test, two-sided). (C) Silencing of *GAPCs* reduces RSV accumulation level. Viral symptoms in RSV infected control plants (TRV:00) and *GAPCs*-silenced plants (TRV:GAPCs). Plants were photographed under normal light at 25 dpi of RSV infection. (D) Relative levels of RSV CP RNA and protein in RSV infected GAPCs-silenced and control plants at 25 dpi, as detected by northern and western blotting, respectively. The RSV CP RNA and CP levels were normalized in relation to rRNA and rubisco, respectively, and the relative levels calculated in relation to the leftmost TRV:00 sample. (E) Silencing of *ATG7* increases RSV accumulation level. Viral symptoms in RSV infected control plants (TRV:00) and *ATG7*-silenced plants (TRV:ATG7). Plants were photographed under normal light at 25 dpi of RSV infection. (F) Relative levels of RSV RNA and CP in RSV infected *ATG7*-silenced and control plants at 25 dpi, as detected by Northern and Western blotting, respectively. The RSV CP RNA and CP levels were normalized in relation to rRNA and rubisco, respectively, and the relative levels calculated in relation to the leftmost TRV:00 sample.

### NbP3IP participates in the regulation of autophagy

We next devised a series of experiments to examine the role of NbP3IP in the absence of RSV p3, once again using transient expression of CFP-NbATG8f to visualize autophagic activity (Han et al., 2015; Huang and Liu, 2015). Thus, we observed an increase of about 2.5-fold in the numbers of autophagosomes following the transient expression of NbP3IP-Myc and CFP-NbATG8f as compared with co-expression of GUSp-Myc and CFP-NbATG8f (Fig. 7A and 7B). Transmission electron microscopy (TEM) was also used to verify the autophagy activation in this study. Compared to the control plants, we could clearly observe increased numbers of autophagic structures in leaves with transient expression of NbP3IP (Fig 7C). There was about a 2-fold increase in the number of visible structures typical of autophagosomes in the cytoplasm (Fig 7D). However, expressing p3 alone could not induce autophagy according to confocal microscopy and TEM observations (Supplementary Fig. S6), In addition, we employed expression of CFP-NbATG8f to monitor autophagy in NbP3IP-silenced TRV VIGS plants (Supplementary Fig. S7A). Confocal microscopy showed that there were many fewer autophagosomes as represented by CFP-NbATG8f puncta in NbP3IP-silenced plants (TRV:NbP3IP) compared with control plants (Fig. 7E, 7F). Compared to TRV:00 control plants, the mRNA levels of ATGs were down-regulated in NbP3IP-silenced plants (Supplementary Fig. S7B). Taken together, these data suggest that NbP3IP is a new player in the regulation of autophagy.

**Fig.7.**
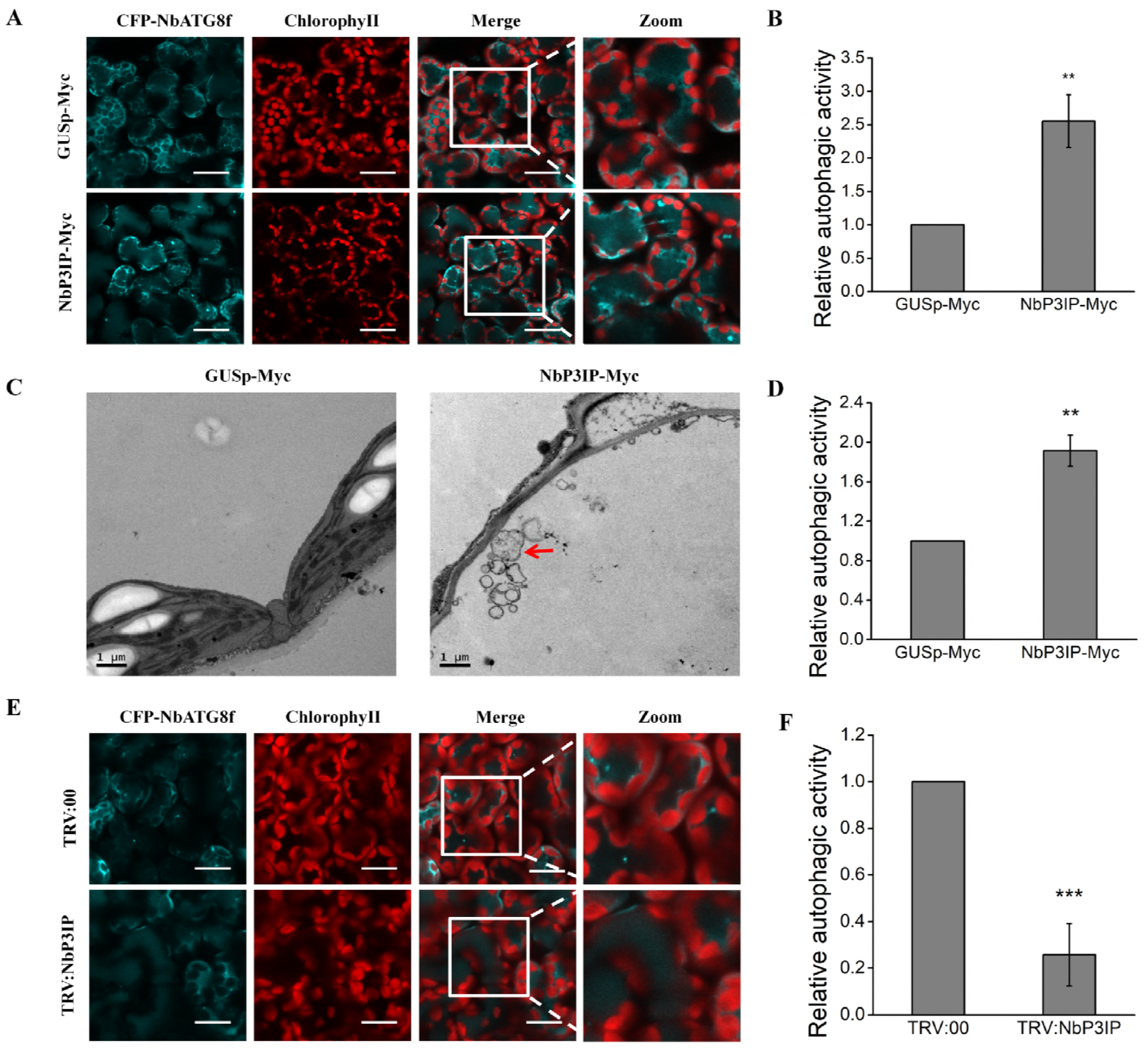
NbP3IP overexpression activates autophagy. (A) Co-expression of NbP3IP-Myc with CFP-NbATG8f increases the appearance of autophagosomes and autophagic vesicles compared to expression of CFP-NbATG8f with a control protein (GUSp-Myc). Images were collected at 60 hpi. Bars, 25 μm. (B) Quantification of increase in autophagic activity in cells imaged in panel A. The autophagic activity was calculated in relation to GUSp-Myc-treated plants. Autophagic bodies were counted from approximately 150 cells for each treatment in three independent experiments. Values represent the mean ± SD. Double asterisks indicates *P*<0.01 of significant difference between GUSp-Myc and NbP3IP-Myc treatments (Student’s *t*-test, two-sided). (C) Examination of autophagic vesicle production by TEM of leaf cells from plants infiltrated with GUSp-Myc or NbP3IP-Myc. Samples collected for processing at 60 hpi. Typical autophagic structures are indicated with red arrows. Bars, 1 μm. (D) Quantification of autophagic vesicles from approximately 20 cells present in TEM images. The autophagic activity is calculated relative to GUSp-Myc-treated plants. The value represents the mean ± SD from three independent experiments. Double asterisks indicates *P*<0.01 of significant difference between GUSp-Myc and NbP3IP-Myc treatments (Student’s *t*-test, two-sided). (E) Silencing of the endogenous *NbP3IP* gene reduces the number of autophagosomes. Confocal images showing *NbP3IP*-silenced (TRV:NbP3IP) or control (TRV:00) plants transiently expressing CFP-NbATG8f at 60 hpi. (F) Relative autophagic activity in *NbP3IP*-silenced plants. The autophagic activity was calculated in comparison to TRV:00 plants. Values represent the mean ± SD from three independent experiments. Approximately 150 cells were used to quantify autophagic structures in each treatment. Three asterisks indicates *P*<0.001 of significant difference between TRV:00 and TRV:NbP3IP treatments (Student’s *t*-test, two-sided).

### RSV infection is reduced in transgenic *N. benthamiana* plants over-expressing NbP3IP

Three transgenic *N. benthamiana* lines (OENbP3IP-1, OENbP3IP-2, OENbP3IP-3) were created in which NbP3IP mRNA levels were increased 3-fold compared to non-transgenic plants (Fig 8A). mRNA levels of autophagy-related genes were similarly increased in the NbP3IP-overexpressing transgenic plants (Fig. 8B). When challenged with RSV, these plants showed delayed and milder symptoms compared to control plants (Fig. 8C and 8D). In addition, the accumulation levels of RSV capsid protein and RNA were reduced at 15 dpi in the systemically infected leaves of the transgenic lines compared with control plants (Fig. 8E). Taken together, our results demonstrate that overexpression of NbP3IP reduces RSV infection in *N. benthamiana* plants.

**Fig.8.**
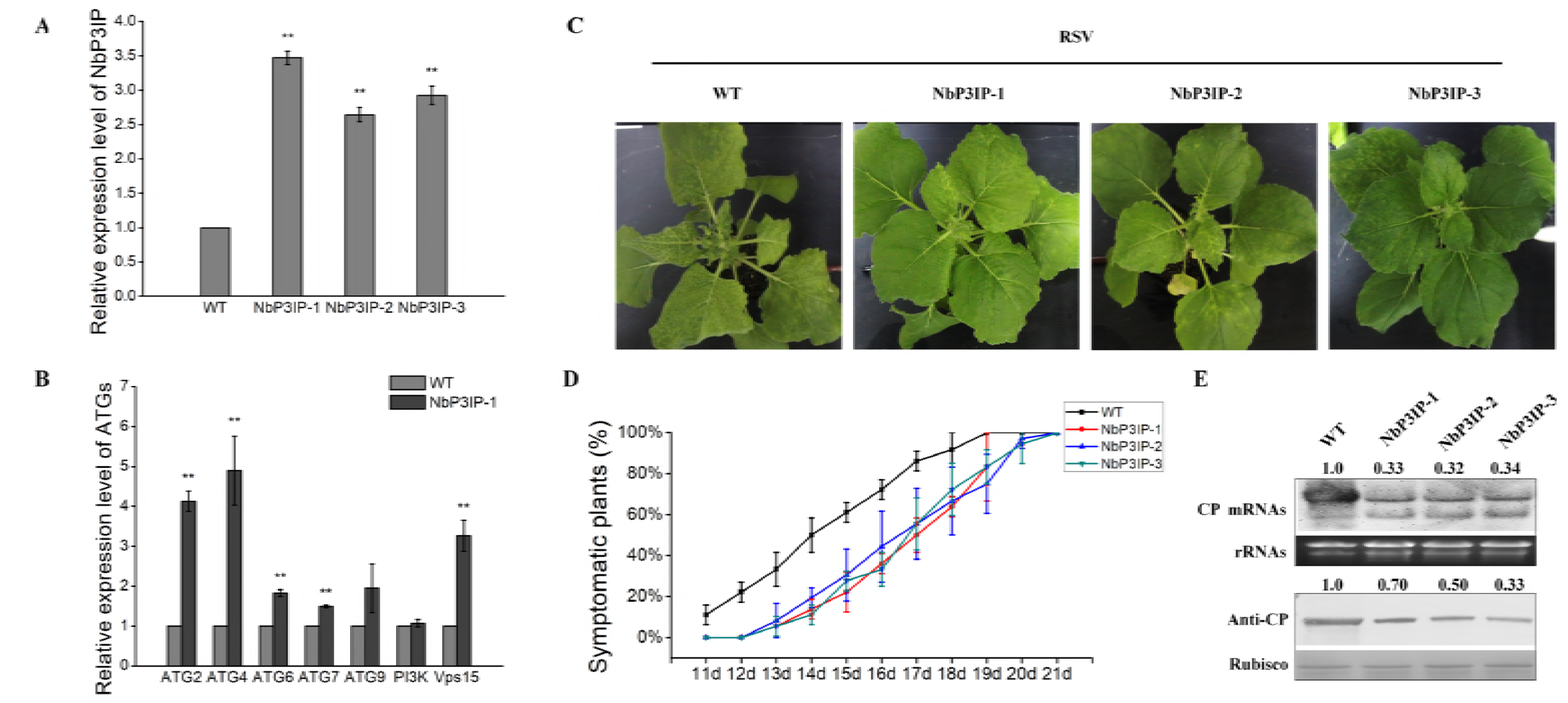
Overexpression of NbP3IP reduces RSV accumulation in infected plants. (A) Expression level of *NbP3IP* in three different transgenic lines (NbP3IP-1, NbP3IP-2 and NbP3IP-3) was analyzed by qRT-PCR with primers listed in supplementary table 1. Values represent means ± SD from three independent experiments. Double asterisks indicate *P*<0.01 of significant difference between wild-type (WT) plants and NbP3IP overexpressed transgenic plants (Student’s *t*-test, two-sided). (B) The expression of ATGs are up-regulated in *NbP3IP* overexpressing transgenic plants. Relative expression of ATGs was monitored by qRT-PCR with primers listed in the supplementary table 1. Values represent means ± SD from three independent experiments. Double asterisks indicate *P*<0.01 of significant difference between WT plants and NbP3IP-1 plants (Student’s *t*-test, two-sided). (C) Overexpression of NbP3IP in transgenic plants alleviates RSV infection. Viral symptoms in plants 15 days post inoculation with RSV. (D) Time-course analysis of RSV infection. Fifteen individual plants each of WT, NbP3IP-1, NbP3IP-2 and NbP3IP-3 transgenic lines were inoculated with RSV and examined daily for appearance of viral symptoms in systemic leaves. Values represent means ± SD from three independent experiments. (E) RSV CP RNA and protein accumulation levels in systemically infected leaves (15 dpi) of WT and NbP3IP-transgenic lines detected by Northern and Western blotting. The value represents RSV CP RNA and protein accumulation normalized to rRNA and rubisco levels, respectively, and expressed relative to the WT plant.

## DISCUSSION

To combat infecting viruses, plants possess several defense mechanisms targeting either the viral nucleic acid or viral proteins. Autophagy, which functions as a pivotal component of host immunity, directly or indirectly retards viral infection through degrading viral proteins. The autophagy-related proteins, NBR1, ATG6 (Beclin-1) and ATG8f, that function as cargo receptor and adaptor proteins, directly interact with virus-encoded proteins and mediate their degradation (Hafren et al., 2017; Haxim et al., 2017; Li et al., 2018). Similarly, a host calmodulin-like protein, rgs-CaM was shown to bind and send the cucumber mosaic virus (CMV) 2b silencing suppressor protein to autolysosomes for proteolytic degradation (Nakahara et al., 2012).

Indeed, the targeting of viral suppressors of RNA silencing (VSRs) is a frequently adopted strategy employed by plants to defend themselves against virus infection. Host factors such as ALY proteins, rgs-CaM (from *Nicotiana tabacum*) and ZmVDE (violaxanthin deepoxidase protein of Zea mays) have all been shown to interfere with the RNA silencing suppression activity of different VSRs (Canto et al., 2006; Nakahara et al., 2012; Chen et al., 2017). In this work, a plant protein (designated as NbP3IP), with a previously unknown function, was identified by screening for interaction with the RSV p3 VSR protein. Using a transient assay, we found that expression of NbP3IP inhibited the VSR ability of p3 by targeting p3 for degradation via the autophagy pathway. Degradation of p3 required the interaction between NbP3IP and p3, and also an interaction between NbP3IP and NbATG8f, a protein known to be involved in autophagosome formation (Nakatogawa et al., 2007; Xie et al., 2008). Furthermore, activation of autophagy is caused by up-regulated expression of NbP3IP, independent of the expression of p3.

In plants, autophagy is induced by environmental stresses, including abiotic stress, such as heat, cold, drought, oxidation, salt, starvation and biotic stress, such as, pathogen invasion and herbivory (Bassham, 2007; Han et al., 2011; Han et al., 2015; Zhu, 2016; Haxim et al., 2017; Hofius et al., 2017; Avin-Wittenberg, 2018; Li et al., 2018). Initiation of autophagy in plants is usually associated with increasing mRNA levels of *ATG* genes, altered accumulation of phytohormones or accumulation of ROS (reactive oxygen species) (Yoshimoto et al., 2009; Perez-Perez et al., 2010; Han et al., 2015; Zhai et al., 2016; Haxim et al., 2017). In starvation-induced mammalian cells, glyceraldehyde 3-phosphate dehydrogenase (GAPDH), phosphorylated by AMP-activated protein kinase (AMPK), redistributes into the nucleus and directly interacts with Sirtuin 1 (Sir1) to cause it to become activated, which is required to initiate autophagy (Chang et al., 2015). Furthermore, Cong Yi *et al*. found that a histone acetyltransferase, which acetylated the K19 and K48 of ATG3, was required for initiation of autophagy in *Saccharomyces cerevisiae* (Yi et al., 2012). The direct mechanism(s) for initiating autophagy in plants, including plant virus-induced autophagy, is largely unknown. Some components of autophagy such as NbATG6 and NbATG8f have been shown to directly participate in plant defense against TuMV and CLCuMuV, respectively (Haxim et al., 2017; Li et al., 2018). In our work, autophagy was activated by the infection of RSV, which also up-regulated the transcription levels of *NbP3IP* and other known autophagy-related genes including *NbATG2*. Similarly, over-expression of NbP3IP in stably transformed plants also increased the expression of *NbATG2* and other autophagy-related genes.

In our study, NbP3IP interacted with NbATG8 to antagonize RSV infection by degrading its VSR p3 protein. However, NbATG8f could not directly interact with the p3 protein, which suggested that NbP3IP might act as an adaptor to form a complex with NbATG8f and p3. Interestingly, protein BLAST analysis identified proteins of unknown function from Arabidopsis, rice and garlic containing small regions of similar sequence (Supplementary Fig. S8). Indeed,one such conserved sequence is the motif YxxL/I (Supplementary Fig. S8), which resembles the LC3-interacting region (LIR) motif found in receptor proteins involved in the formation of phagosomes during selective autophagy processes (Birgisdottir et al., 2013). Hence, we argue that the involvement of P3IP with autophagy might exist and be conserved in other plant species.

Taken together, our results demonstrate that NbP3IP functions as a newly identified regulator of autophagy to regulate viral infection in plants and that autophagy also contributes to plant antiviral defense against the negative-strand RNA virus RSV. Based on our results, a working model of NbP3IP-mediated autophagy defense against RSV infection can be outlined (Fig. 9), whereby, RSV infection induces the expression of NbP3IP, which activates autophagy. NbP3IP recognizes RSV p3 and carries it to the autophagosome by interacting with NbATG8f. Degradation of p3 prevents it from interfering with the plant RNA silencing system, thereby allowing the RSV RNAs to be targeted and RSV infection to be constrained

**Fig.9.**
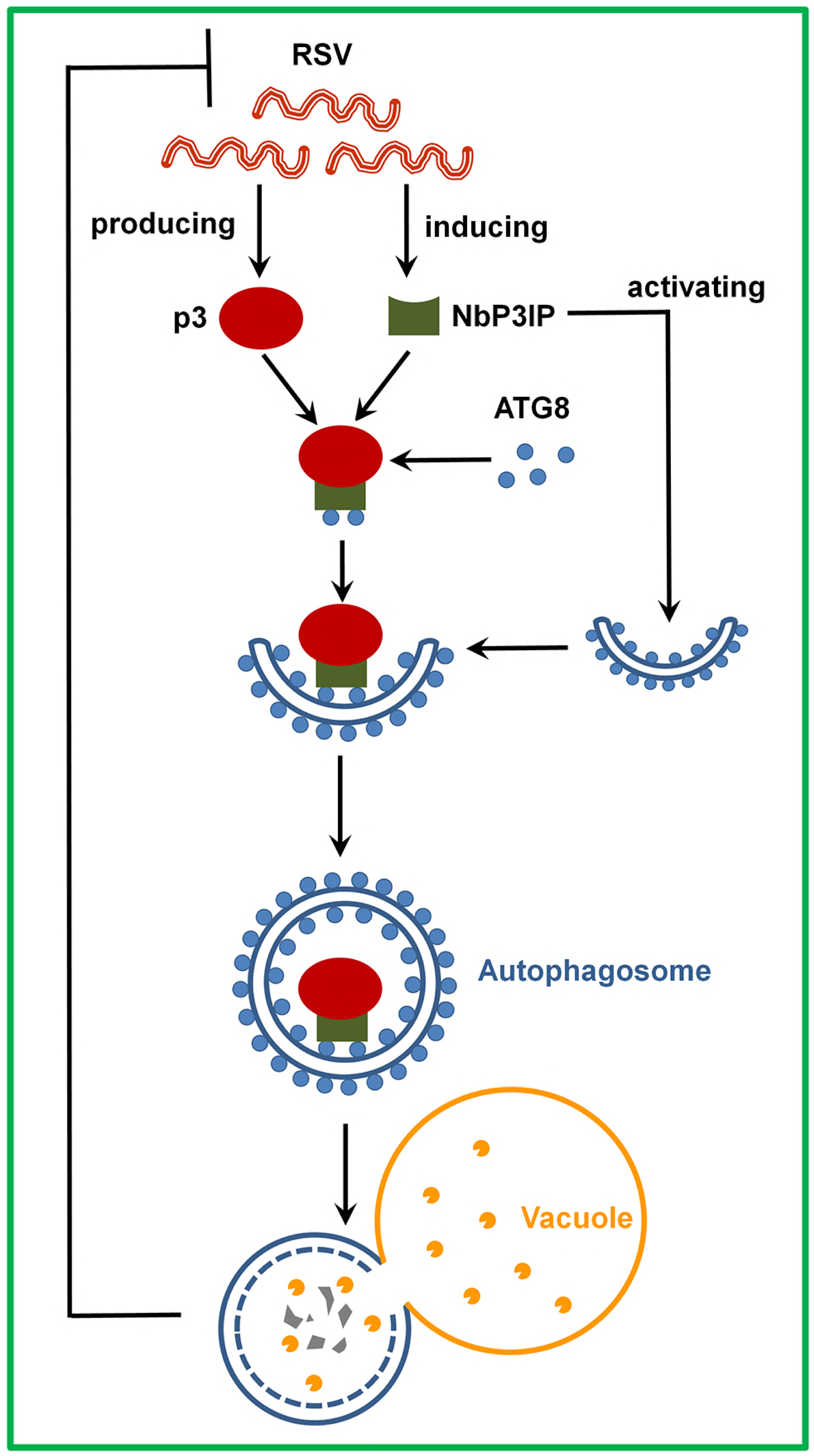
Working model for the role of NbP3IP in stimulating autophagy to resist RSV infection. As the RNA silencing suppressor protein of RSV, p3 is produced in RSV infected plants, however, the infection of RSV also induces the expression of NbP3IP, which interacts with p3 to activate autophagy. NbP3IP interacts with p3 and carries p3 to the autophagosome by cooperating with NbAtg8f that is a key factor in autophagy. The removal of p3 via the autophagy pathway enhances the activity of the plant RNA silencing system in targeting the RSV RNAs for degradation.

## METHODS

### Plant materials and growth conditions

GFP-transgenic line 16c *N. benthamiana*, wild type *N. benthamiana* and OENbP3IP-transgenic *N. benthamiana* plants were grown in pots under constant conditions of 60% relative humidity and a 16h light/8h dark photoperiod. For transgenic overexpression of *NbP3IP* which was cloned into the pCV vector (Lu et al., 2011), Agrobacterium-mediated transformation of *N. benthamiana* plants was conducted following a standard protocol (Horsch RB, 1985), and the regenerated transformants were screened as previously described (Shi et al., 2016).

### Plasmid construction

Gene sequences were amplified by PCR using *ExTaq* DNA Polymerase (TaKaRa) for cloning purposes (primers are listed in supplementary table 1). The full length clones of RSV p3 and pc3 were generated from cDNA derived from *Oryza sativa* infected with Rice stripe virus and the full length clone of NbP3IP was generated from cDNA derived from *N. benthamiana*. For the Y2H assay, genes were ligated into pGBKT7 and pGADT7 to yield pGBK-NbP3IP, pGBK-NbP3IP-S1, pGBK-NbP3IP-S2 and pGAD-p3, respectively. For BiFC assays, inserts were ligated into pCV-YFPn-C and pCV-YFPc-C (YFP added as an N-terminal fusion) (Lu et al., 2011) to construct pCV-YFPn-NbP3IP, pCV-YFPc-NbP3IP, pCV-YFPn-p3, pCV-YFPc-p3, pCV-YFPn-NbP3IP-S2, pCV-YFPn-NbATG8f and pCV-YFPc-NbATG8f. Plasmids pCV-YFPn-GUS, pCV-YFPc-GUS were described previously (Jiang et al., 2014). For transient expression analysis in plants cells, full length NbP3IP, p3 and pc3, and the partial fragments of NbP3IP (NbP3IP-S1, NbP3IP-S2), flanked with *Xba*I and *Kpn*I restriction sites, were introduced into vector pCV-eGFP (Lu et al., 2011) for subcellular localization studies; the full length of NbP3IP, p3 and pc3 with a C-terminal Myc tag, the partial fragment of NbP3IP (NbP3IP-S2) and GUS (GUSp) with C-terminal tagged Myc, flanked with *Xba*I and *Sac*I restriction sites, were introduced into pCV empty vector for transient overexpression experiments (Lu et al., 2011). The full length clone of NbATG8f was generated from cDNA derived from *N. benthamiana*. The resulting DNA fragment was purified and transferred into the entry vector pDONR207 (Invitrogen) by recombination using BP Clonase (Invitrogen), pDONR207 clones were further transferred into the Gateway vector pGWB6 (Nakagawa et al., 2007) (GFP as an N-terminal fusion) to yield pGWB6-NbATG8f.

### Virus-induced gene silencing

A partial sequence of NbP3IP was amplified with primers that are listed in supplementary table 1 and cloned into pTRV2 using flanking *Cla*I and *Sal*I restriction sites, producing the vector TRV:NbP3IP. Viral infection by Agrobacterium infiltration to initiate NbP3IP silencing was performed as described previously (Peng et al., 2011).

### RNA extraction and Northern blotting

Total RNA was extracted from plants using Trizol (Invitrogen, Carlsbad, California, USA) according to the manufacturer’s instructions. For Northern blot analyses, DNA templates for synthesis of RSV CP and mGFP probes were amplified with primers that are listed in supplementary table 1. The probes were labeled with digoxigenin (DIG) according to the manufacturer’s protocol (DIG High Prime DNA Labeling and Detection Starter Kit II, Roche, Basel, Switzerland). Northern blot procedures were performed as previously described (Jiang et al., 2014). For small RNA Northern blotting, 20 μg of total RNA was separated on a 15% polyacrylamide gel, and transferred electrophoretically to Amersham Hybond™-NX membranes (GE Healthcare) using H_2_O for 1h (Guo et al., 2012). Chemical crosslinking was conducted to fix the RNA to the membrane (Pall et al., 2007). Four different segments of the GFP gene (150bp) were used as templates to synthesis DIG-labeled probes according to the manufacturer’s protocol (DIG High Prime DNA Labeling and Detection Starter Kit II, Roche, Basel, Switzerland). ULTRAhyb®-Oligo Hybridization Buffer (Ambion) was used to optimize the hybridization of probes to siRNAs in our Northern analysis. The software Image J was applied to do quantitative calculations of digital images of Northern blots.

### qRT-PCR, semi-quantitative RT-PCR analysis

The genomic DNA was removed from purified total RNA by RNase-free DNase I treatment before qRT-PCR and semi-quantitative RT-PCR (gDNA wiper, Vazyme). The cDNA was synthesized according to the manufacturer’s protocol (HiScript® II Q RT SuperMix for qPCR (+gDNA wiper), Vazyme). qRT-PCR was used to measure the expression of NbP3IP using primers that are listed in supplementary table 1, and to confirm the silencing of ATGs using the primers described previously (Wang et al., 2013). The *N. benthamiana* Ubiquitin C (UBC) gene (Accession Number: AB026056.1) was used as the internal reference gene for analysis and the primers are listed in supplementary table 1 (Rotenberg et al., 2006; Shi et al., 2016). A Roche Light Cycler®480 Real-Time PCR System was used for the reaction and the results were analyzed by the ΔΔC_T_ method. Semi-quantitative RT-PCR was used to measure the expression of p3-GFP and free GFP at 26 cycles using primers listed in supplementary table 1 (Peng et al., 2011).

### Y2H, BiFC and analysis of subcellular localization

The Matchmaker Yeast Two-Hybrid System 3 (Clontech) was used for yeast two-hybrid assays to examine the p3 interaction with NbP3IP. The experiments were performed as described previously (Shi et al., 2007). The vectors used in BiFC and subcellular localization assay experiments have been described previously (Lu et al., 2011; Yan et al., 2012). These assays were performed using laser scanning confocal microscopy as described previously (Jiang et al., 2014).

### Co-immunoprecipitation (Co-IP) assay

For Co-IP assays, the proteins under study were transiently expressed by agro-infiltration in *N. benthamiama* plants. At 3 dpi, leaf samples (8 discs) were extracted in ice-cold extraction buffer (GTEN buffer (10% glycerol, 25 mM Tris-HCl, pH 7.5, 1 mM EDTA, and 150 mM NaCl), 10 mM DTT, 1 mM PMSF, 0.15% Nonidet P40, and 1×protease inhibitor cocktail (Roche) (Wang et al., 2015). Leaf lysates were then mixed with anti-GFP mAb-Magnetic beads (MBL, Japan), mixed well and incubated with gentle agitation for an hour at room temperature. The beads were collected by magnet and rinsed 3 times with washing buffer (GTEN butter, 1 mM PMSF and 1×protease inhibitor cocktail, freshly prepared), and then the bound complexes were eluted by boiling with protein loading buffer for 5 minutes. The samples were separated by SDS-PAGE in a 12% polyacrylamide gel and detected using an anti-Myc antibody (Han et al., 2015).

### Western blotting

Total proteins of plant leaf samples were extracted with lysis buffer (100 mM Tris-HCl, pH 8.8, 60% SDS, 2% β-mercaptoethanol). Proteins were fractionated by 12% SDS-PAGE, transferred onto nitrocellulose membrane (Amersham, Germany), then detected using anti-GFP polyclonal primary antiserum and alkaline phosphatase-conjugated anti-mouse (TransGen) secondary antibody. The antigen–antibody complexes were visualized using nitrotetrazolium blue chloride/5-bromo-4-chloro-3-indolyl phosphate (NBT/BCIP) buffer (Sigma-Aldrich, St. Louis, Missouri, USA) under standard conditions. Image J software was used to do quantitative calculations of digital images of Western blots.

### Leaf chemical treatment, confocal microscopy and TEM

Phosphate-buffered saline containing 2% dimethyl sulfoxide (DMSO, as control) or an equal volume of DMSO with 100 μM MG132 (Sigma) for inhibition of the 26S proteasome, or H_2_O as a control and an equal volume of H_2_O containing 10 mM 3-MA (Sigma) for inhibition of autophagy, was infiltrated into leaves 16 h before samples were collected. Confocal imaging was performed as described (Jiang et al., 2014). Alternatively, the leaves were agroinfiltrated with the autophagy marker CFP-NbATG8f for a 60 h expression period, followed by additional infiltration with 20 μM E-64d (Sigma) for an 8 h period. A Leica TCS SP5 (Leica Microsystems, Bannockburn, IL, USA) confocal laser scanning microscope was used to examine the fluorescence of CFP with an excitation light of 405 nm, and the emission was captured at 454 to 581 nm.

For TEM observations, the 20 μM E-64d-treated leaves were cut into small pieces (2×2 mm^2^). The treatments and the examination of the sampled tissues were performed as described before (Li et al., 2017).

## ACKNOWLEDGEMENTS

This work was supported by the State Basic Research Program of China (2014CB138403), the national key research and development program of China (2017YFA0503401), the National Natural Science Foundation of China (31772239), the Major Project of New Varieties of Genetically Modified Organism of China (2016ZX08001-002), the Rural & Environment Science & Analytical Services Division of the Scottish Government, the International Science & Technology Cooperation Program of China (2015DFA30700) and the K.C. Wong Magna Fund of Ningbo University.

## AUTHOR CONTRIBUTIONS

L.L.J., Y.W.L., X.Y.Z., F.Y., Y.L. and J.C. designed experiments and analyzed data. L.L.J., Y.W.L., X.Y.Z., X.Y., Y.C., T.Z., XING Z., S.W., X.Z., X.X.Z., J.P., H.Z. and L.L. performed the experiments. L.L.J., Y.W.L., X.Y.Z., F.Y., Y.L. and J.C. prepared the figures. X.S. provided the help with TEM experiment. S.M.F. provided constructive comments and help with writing. L.L.J., Y.W.L, X.Y.Z., F.Y., Y.L. and J.C. wrote the paper.

## Supplementary Figure

**Supplementary Figure S1.**
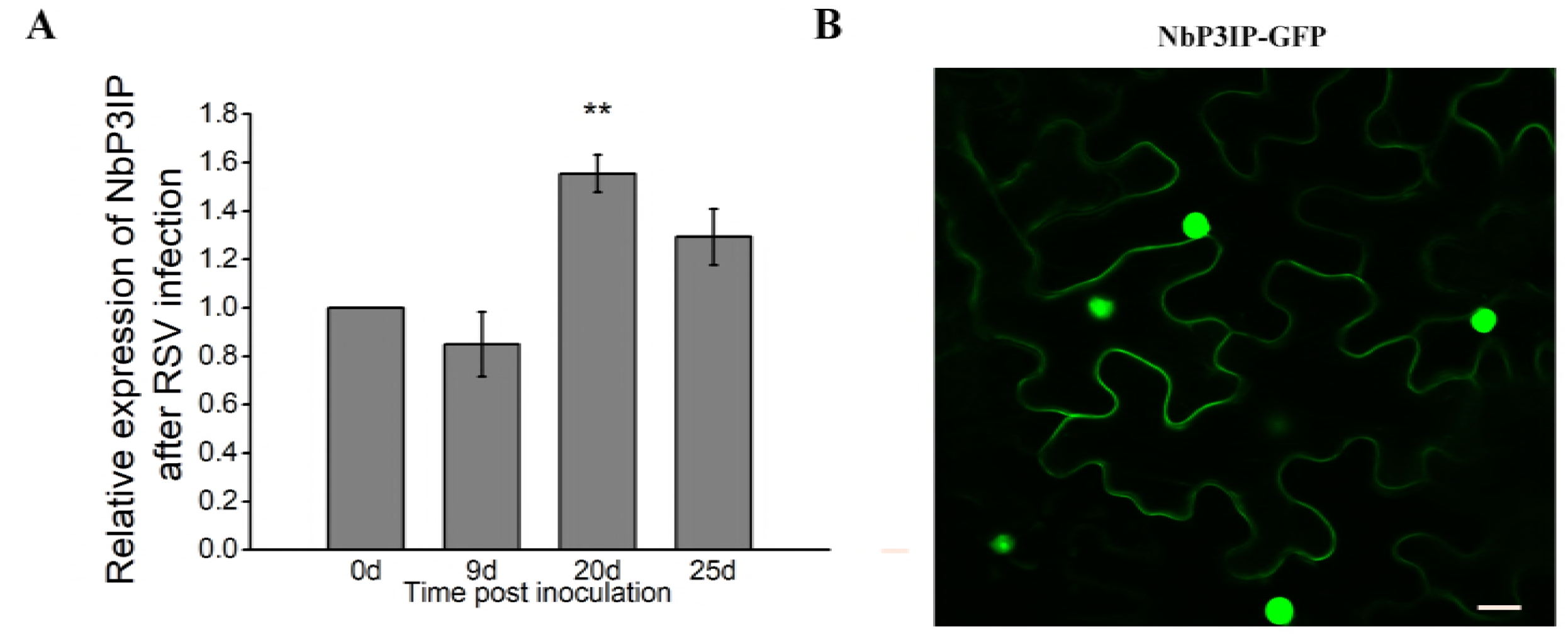
Expression pattern and subcellular localization of NbP3IP. (A) Relative expression level of *NbP3IP* after RSV infection. *NbP3IP* expression was monitored by qRT-PCR at 0, 9, 20, 25 dpi. Values represent means ± SD from three independent experiments. Double asterisks indicate *P*<0.01 of significant difference between RSV infected 0 day plants and other plants (Student’s *t*-test, two-sided). (B) Subcellular localization of NbP3IP-GFP. Confocal images showing the expression of NbP3IP-GFP in *N. benthamiana* leaf cells. Bars, 50 μm.

**Supplementary Figure S2.**
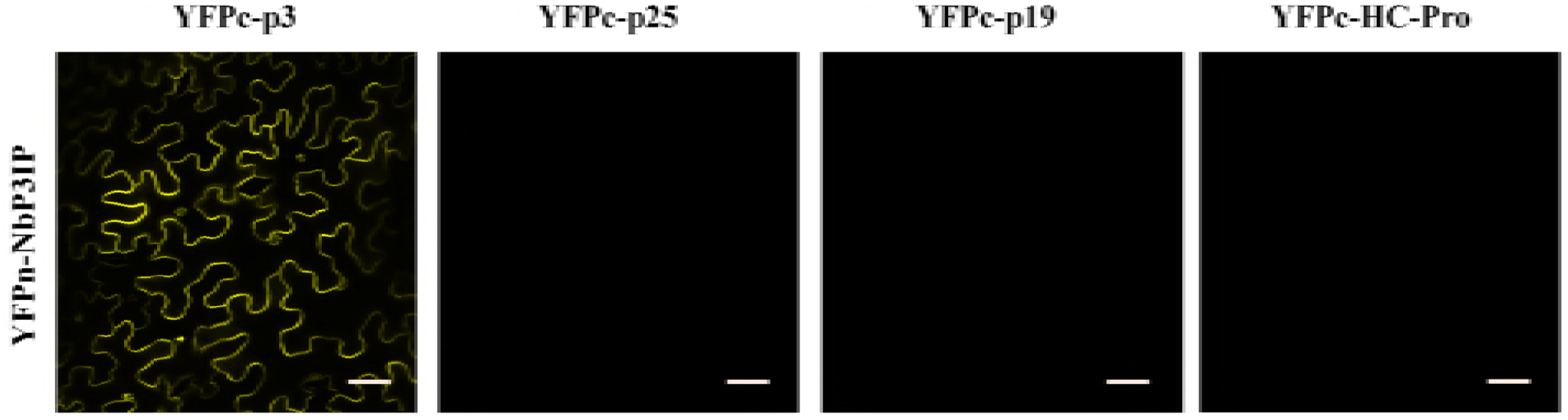
NbP3IP interacts with p3 but not other VSRs. BiFC assays between NbP3IP and RSV p3, PVX p25, TBSV p19, TuMV HC-Pro in the leaves of *N. benthamiana* at 60 hpi. The N-terminal YFP fragment (YFPn) was fused to the N-terminus of NbP3IP, and the C-terminal YFP fragment (YFPc) was fused to N-terminus of p3, p25, p19, HC-Pro. Bars, 50 μm.

**Supplementary Figure S3.**
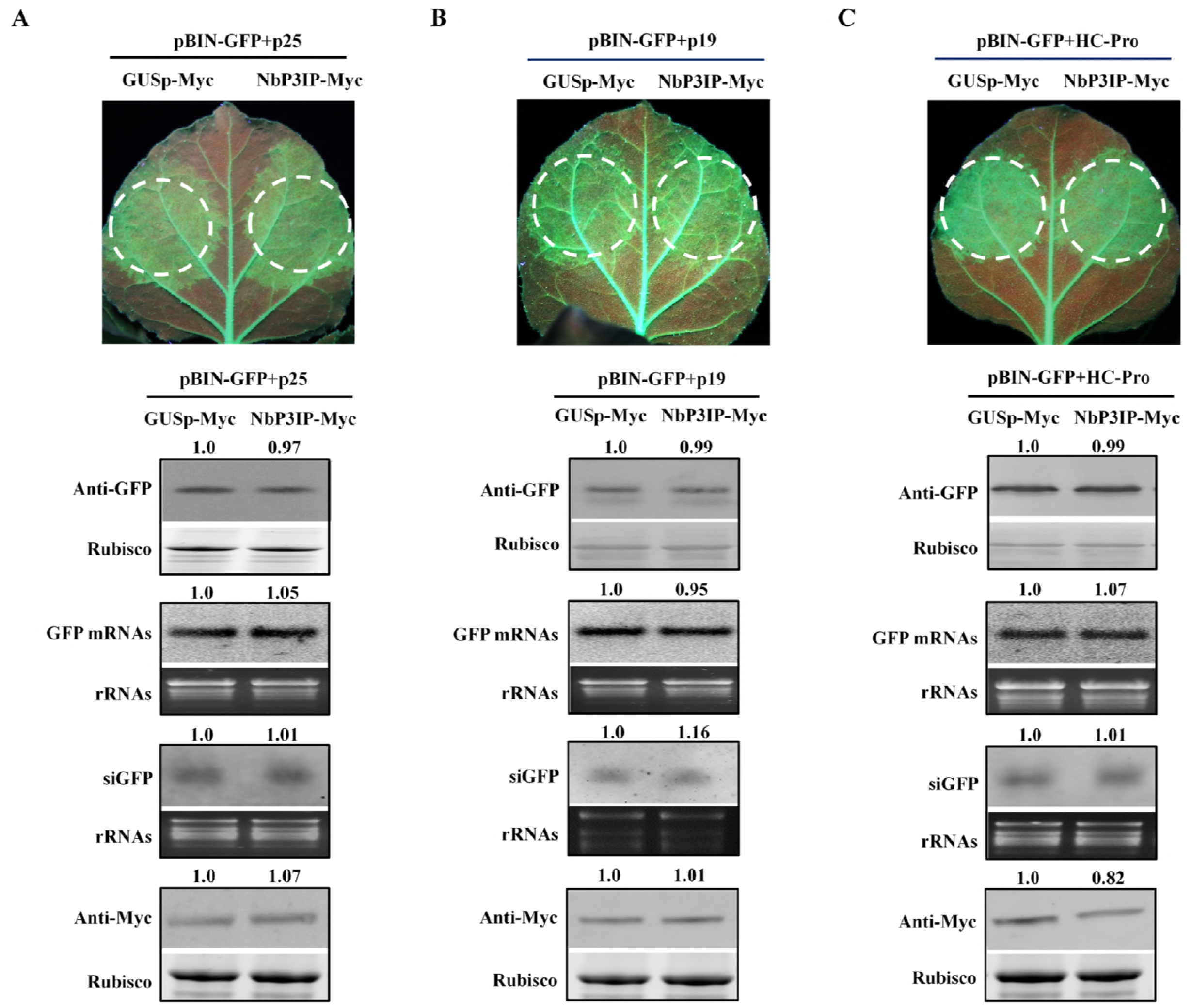
The expression of NbP3IP has no effect on the VSR ability of other suppressors. The expression of NbP3IP did not affect the VSR ability of PVX p25 (Panel A), TBSV p19 (Panel B) or TuMV HC-Pro (Panel C). pBIN-GFP, viral suppressor and NbP3IP (right patch) were co-infiltrated in 16c transgenic *N. benthamiana* plants leaves, with GUSp-Myc (left patch) substitution of NbP3IP-Myc as a control treatment. The patches were photographed under UV lamp at 5 dpi. The accumulations of GFP protein and mRNA were detected by western blot and northern blot. The accumulation of GFP-derived siRNA was analyzed by small RNA northern blot. The expression of GUSp-Myc and NbP3IP-Myc protein were detected by western blot with Myc antibody. The relative proteins levels in were normalized to rubisco, and the GFP mRNA and siRNA levels were normalized to rRNA. Relative values were calculated in comparison to the GUSp-Myc treatment (left patch), and this value was set as standard 1.

**Supplementary Figure S4.**
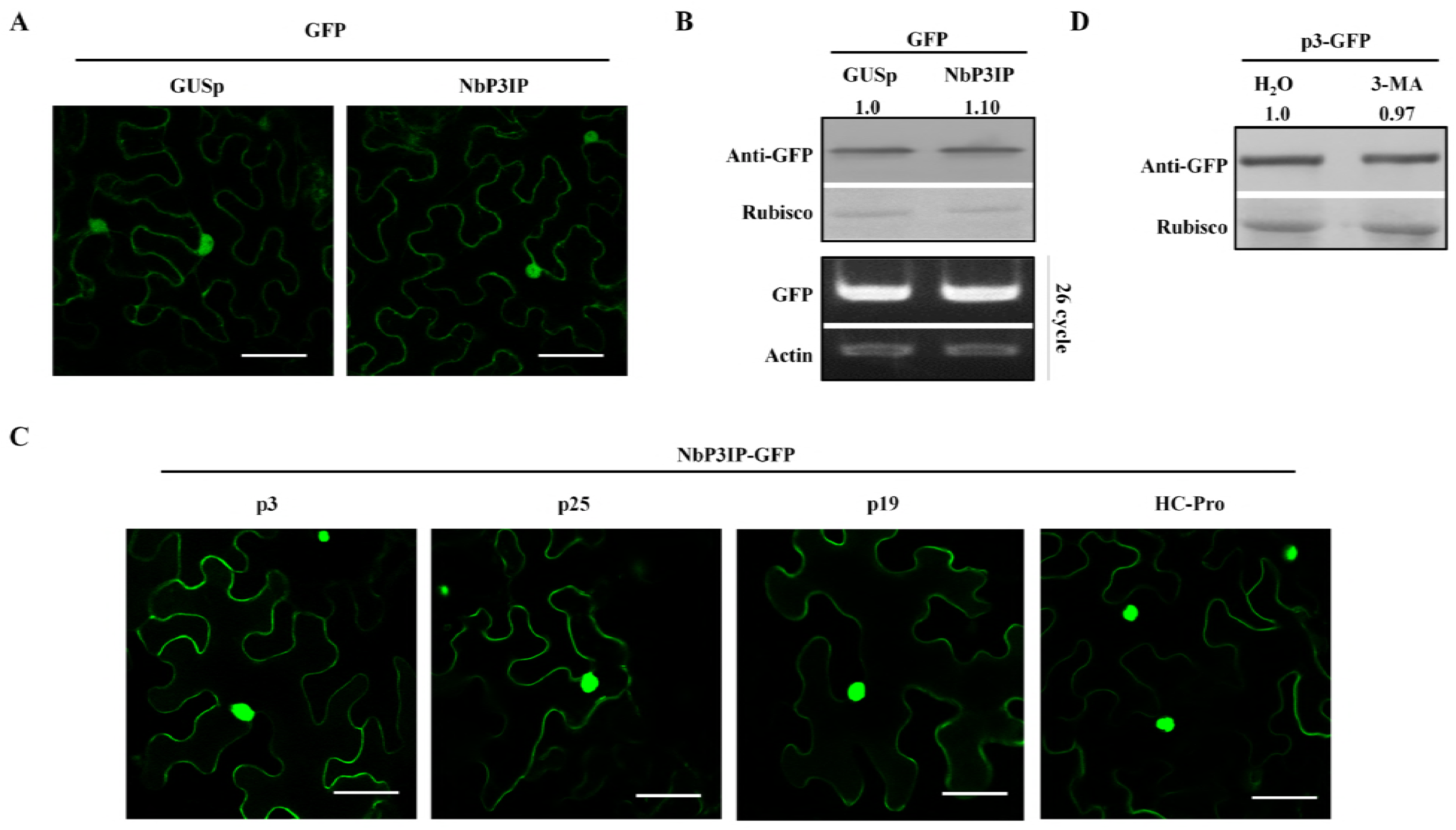
Expression of NbP3IP has no effect on GFP accumulation and the autophagy inhibitor 3-MA has no effect on p3 protein accumulation. (A and B) The accumulation of unfused GFP is not affected by co-expression with NbP3IP. (A) Micrographs show cells co-expressing GFP and NbP3IP (right) or GFP and GUSp (left) as a control. Infiltrated *N. benthamiana* leaves were examined at 60 hpi. Bars, 25 μm. (B) Western blot of total protein extracts from (A) detected with GFP antibody. Semi-quantitative RT-PCR was used for analysis of GFP transcripts. Actin served as an internal sqRT-PCR standard. The GFP accumulation when co-expressed with GUSp was normalized to rubisco, and this value used for comparative calculations. (C) The subcellular localization of NbP3IP is not affected by co-expression of p3 and other VSRs. Images show cells co-expressing NbP3IP-GFP with RSV p3, PVX p25, TBSV p19, TuMV HC-Pro respectively. Infiltrated *N. benthamiana* leaves were examined at 60 hpi. Bars, 25 μm. (D) The autophagy inhibitor 3-MA does not affect accumulation of p3-GFP. Expression p3-GFP in *N. benthamiana* leaves for 48 h, followed by H_2_O or 10 mM 3-MA treatment for 16 h. Proteins were detected by western blotting using an anti-GFP antibody. The value represents p3-GFP protein accumulation relative to rubisco, and the protein accumulation with H_2_0 treatment used for comparison.

**Supplementary Figure S5.**
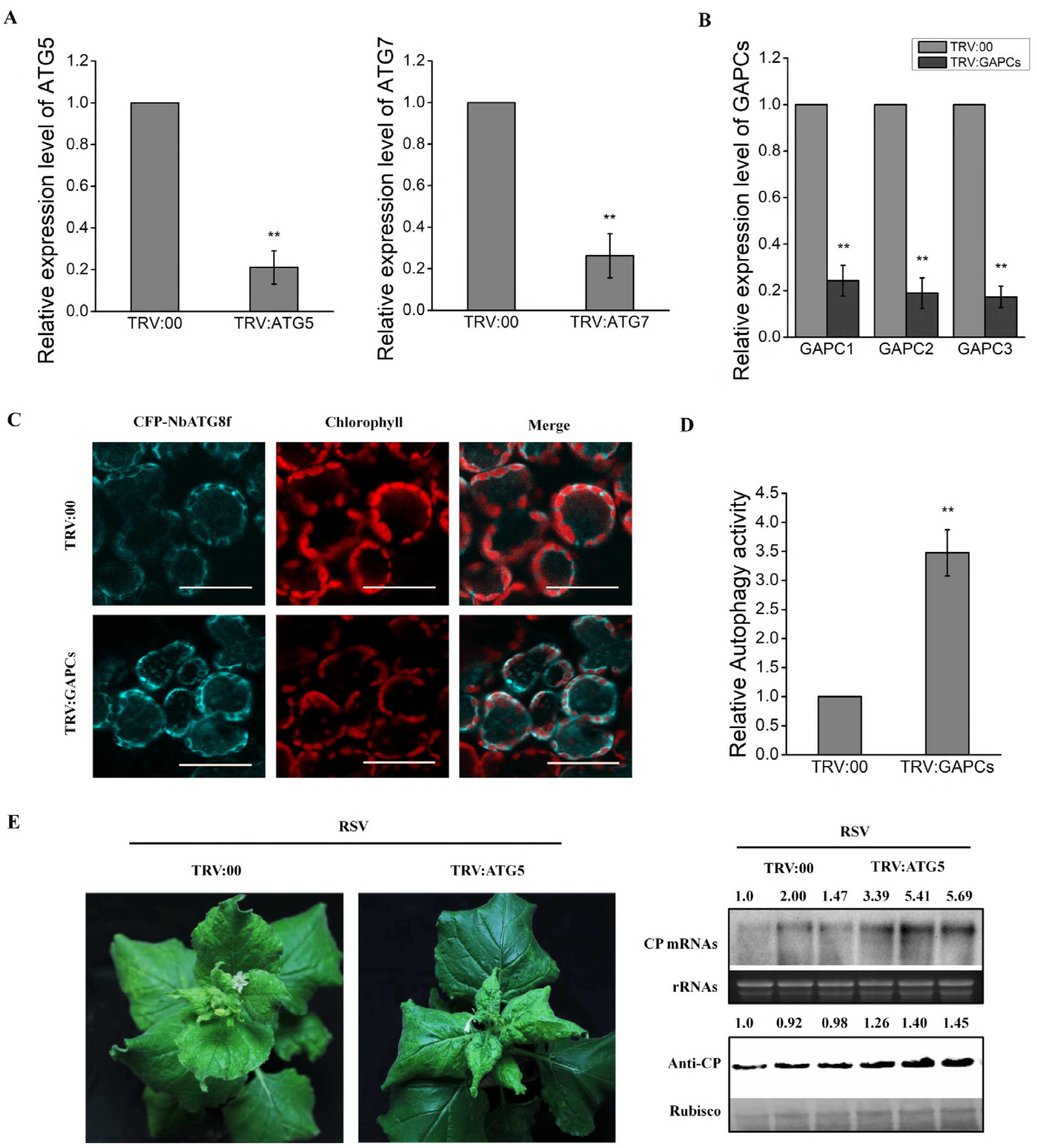
Confirmation of the silencing of ATG5, ATG7 and GAPCs, and that silencing of GAPCs activates autophagy. (A) Real-time RT-PCR analysis to show relative expression levels of ATG5 and ATG7 following TRV VIGS-mediated silencing. Primers are listed in the supplementary table 1. Upper leaves were examined at 15 dpi. Values represent means ± SD from three independent experiments. (each sample containing three biological repeats and each biological repeat consisting of three technical replicates). Double asterisks indicate *P*<0.01 of significant difference between TRV:00 control plants and TRV:ATG5 or TRV:ATG7 silenced plants (Student’s *t*-test, two-sided). (B) Real-time RT-PCR analysis of relative expression levels of three GAPCs after TRV-mediated silencing. Primers are listed in the supplementary table 1. Upper leaves were examined at 15 dpi. Values represent means ± SD from three independent experiments. (each sample containing three biological repeats and each biological repeat consisting of three technical replicates). Double asterisks indicate *P*<0.01 of significant difference between TRV:00 control plants and TRV:GAPCs silenced plants (Student’s *t*-test, two-sided). (C) Silencing of GAPCs actives autophagy. Confocal micrographs showing the autophagy-specific marker CFP-NbATG8f expressed in GAPCs-silenced plants (TRV:GAPCs) and control (TRV:00) plants. Autophagosomes and autophagic bodies are revealed as CFP positive puncta in mesophyll cells. CFP-NbATG8f fusion proteins are in cyan, and chloroplasts are in red. Bars, 25 μm. (D) Relative autophagic activity in GAPCs-silenced plants. The autophagic activity in TRV:00 control plants were used as the comparator. Values represent the mean ±SD from three independent experiments. Approximately 150 cells were used to quantify autophagic structures in each treatment. Double asterisks indicate *P*<0.01 of significant difference between TRV:00 control plants and TRV:GAPCs silenced plants (Student’s *t*-test, two-sided). (E) Silencing of *ATG5* increases RSV accumulation level. Viral symptoms in RSV infected control plants (TRV:00) and *ATG5*-silenced plants (TRV:ATG5). Plants were photographed under normal light at 25 dpi of RSV infection.

**Supplementary Figure S6.**
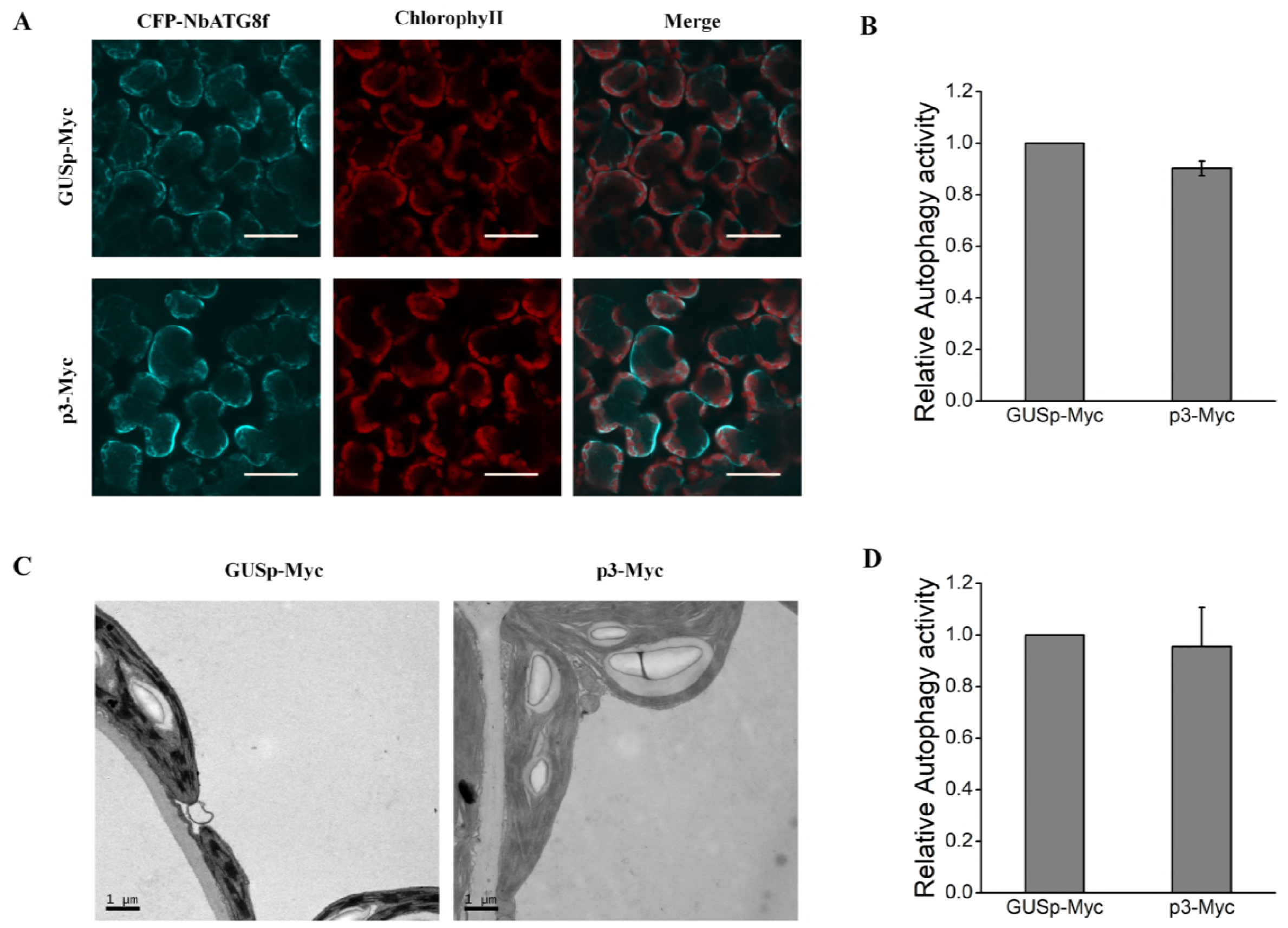
Transient expression of p3 does not induce autophagy. (A) Autophagic body production, as revealed by CFP-NbATG8f-labelled vesicle production, is not induced by p3. CFP-NbATG8f fusion proteins are in cyan, and chloroplasts are in red. Bars, 25μm. (B) Relative autophagic activity in plants transiently expressing GUSp-Myc or p3-Myc. Values represent the mean ±SD from three independent experiments. Approximately 150 cells were used to quantify autophagic structures in each treatment. (C) Representative TEM imagines from *N. benthamiana* leaf cells infiltrated with GUSp-Myc or p3-Myc at 60 hpi. Bars, 1 μm. (D) Relative autophagic activity in plants transiently expressing GUSp-Myc or p3-Myc. Values represent the mean ± SD from three independent experiments. Autophagic bodies were counted from approximately 20 cells in each treatment.

**Supplementary Figure S7.**
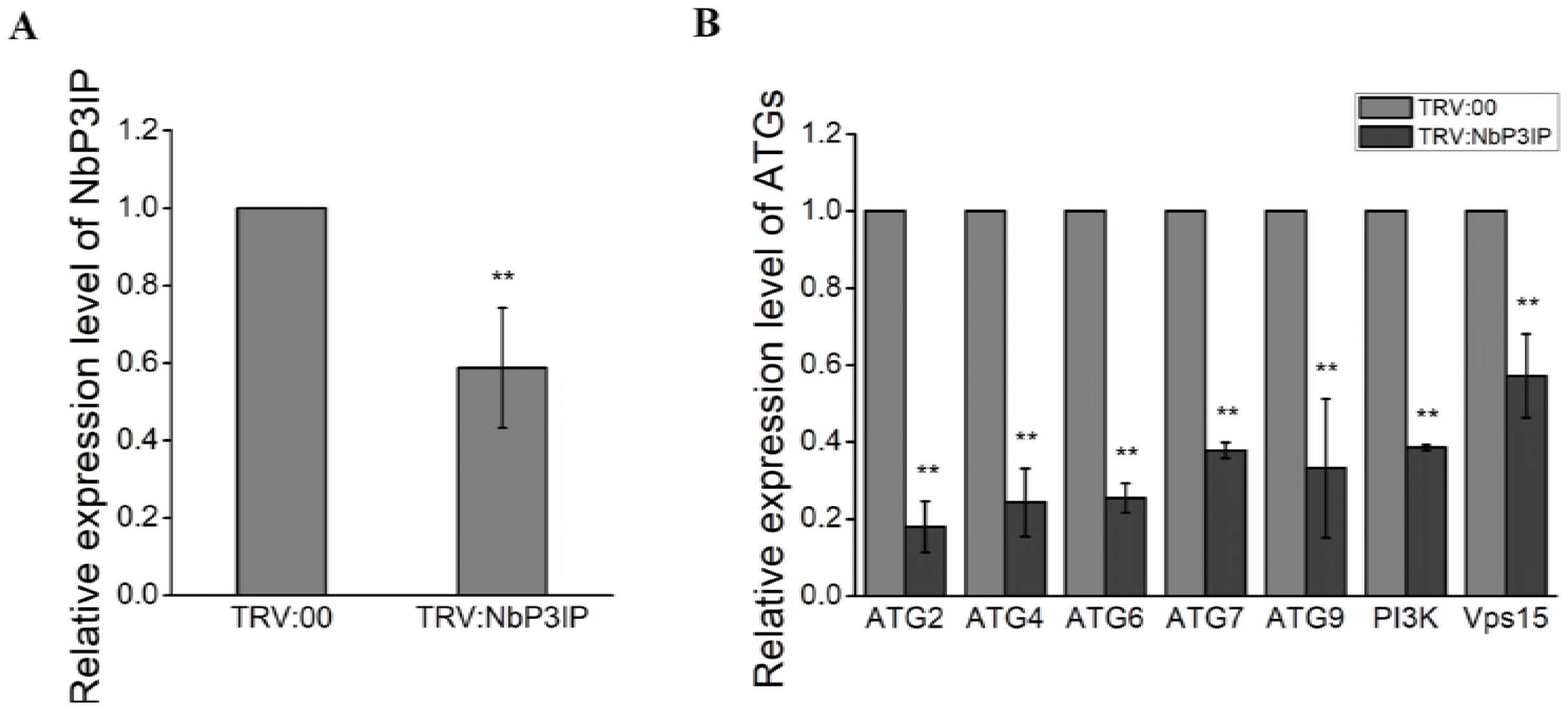
Silencing of *NbP3IP* inhibits the expression of ATGs. (A) Real-time RT-PCR analysis demonstrates the silencing of *NbP3IP*. Systemic leaves were examined after silencing of *NbP3IP* via TRV-based VIGS at 15 dpi. Values represent means ± SD from three independent experiments. (each sample containing three biological repeats and each biological repeat consisting of three technical replicates). Double asterisks indicate *P*<0.01 of significant difference between TRV:00 control plants and TRV:NbP3IP silenced plants. (Student’s *t*-test, two-sided). (B) Real-time RT-PCR analysis of the relative expression of ATGs in TRV:NbP3IP-silenced plants compared to TRV:00 plants. Values represent means ± SD from three independent experiments. (each sample containing three biological repeats and each biological repeat consisting of three technical replicates). Double asterisks indicate *P*<0.01 of significant difference between TRV:00 control plants and TRV:NbP3IP silenced plants. (Student’s *t*-test, two-sided).

**Supplementary Figure S8.**
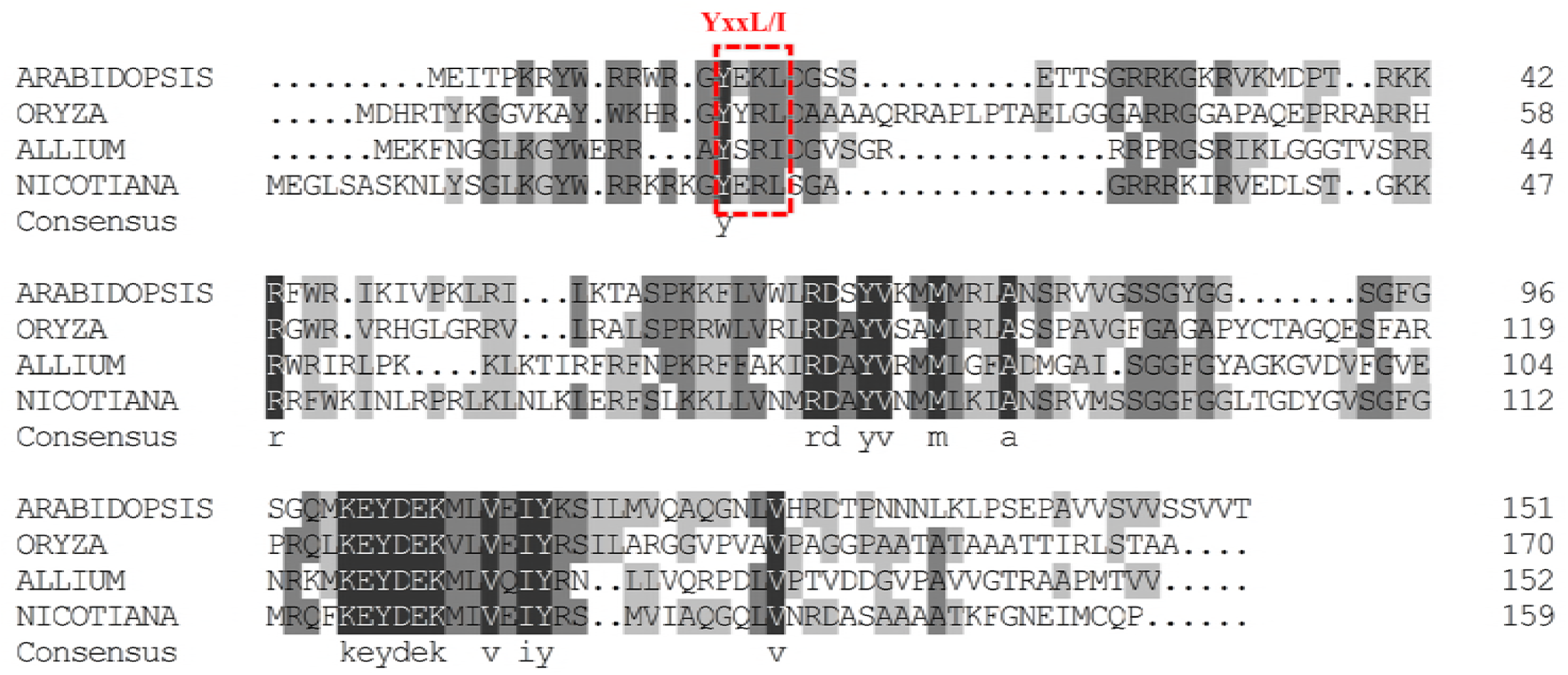
Sequence alignment of NbP3IP in different plant species. Amino acid sequence alignment of NbP3IP with similar proteins from *Arabidopsis thaliana* (NP_191824.1), *Oryza sativa* (XP_015644236.1), *Allium sativum* L (ADK23953.1) and *Nicotiana benthamiana* (Niben101Scf03390g06004.1) reveals a low overall homology among these proteins but identifies three conserved regions including the autophagy-related LIR motif.

Supplementary table S1.

List of Primers Used in This Study

